# Buc2l Drives Higher-Order Germ Plasm Assembly in Zebrafish

**DOI:** 10.64898/2026.11.20.767596

**Authors:** Fiona Carey, Walter Bronkhorst, Martin M. Möckel, Yasmin El Sherif, Miguel A. Andrade-Navarro, René F. Ketting

**Affiliations:** Institute of Molecular Biology, Ackermannweg 4, 55128 Mainz, Germany; International PhD Programme on Gene Regulation, Epigenetics & Genome Stability, 55128 Mainz, Germany; Johannes Gutenberg University, Institute of Developmental Biology and Neurobiology, 55099 Mainz, Germany

## Abstract

Germ plasm formation is essential for primordial germ cell (PGC) specification, yet its underlying molecular mechanisms remain incompletely understood. Here, we identify Buc2l as a novel factor required for extensive germ plasm assembly during zebrafish embryogenesis. While dispensable for Balbiani body formation during oogenesis, Buc2l is essential downstream in the early embryo, where it coalesces key components including Ziwi, Buc, and Tdrd6a/b, into robust assemblies along the cleavage furrows. In *buc2l* mutants, germ plasm becomes fragmented, leading to a marked loss of PGCs, a male-skewed sex ratio, and adult sterility. Unlike the germ plasm component Buc, Buc2l lacks a prion-like domain. Instead, it features a tandem repeat region. *In vitro* and *in vivo* analyses reveal that the Buc2l repeat region stimulates the aggregation of small Buc granules into larger assemblies. Buc2l variants that lack varying numbers of the tandem repeats still incorporate into Buc condensates, yet fail to drive assembly growth. Given that the tandem repeat numbers vary among different species, these findings demonstrate how tandem repeat expansion may tune condensate size, revealing an evolutionary strategy for germ plasm assembly control.

## INTRODUCTION

Germ cell fate in multicellular organisms is established through one of two main mechanisms: induction or preformation (Extavour and Akam, 2003; Whittle and Extavour, 2017). Induction, considered to be the ancestral mode, relies on inductive signalling through the BMP and Wnt pathways, where somatic cells direct neighbouring cells to take on a germline fate, as is observed in mammals (Ying and Zhao, 2001). In contrast, many species with rapid embryonic development such as the zebrafish (*Danio rerio*) and *Caenorhabditis elegans* employ preformation. In this case, maternally deposited cytoplasmic determinants, termed germ plasm, are inherited by a subset of cells in developing blastomeres that will give rise to primordial germ cells (PGCs) (Extavour and Akam, 2003; Knaut et al., 2000; Updike and Strome, 2010). The existence of such a dedicated germline cytoplasm was first proposed by August Weismann (1892), who postulated a distinction between germ and somatic cells. Since then, many studies have refined our understanding of germ plasm; however, Weismann’s fundamental principle of inheritance through the germline remains consistent.

During oogenesis in *D. rerio*, a transient, amyloid-like structure is formed, known as the Balbiani body (Bb) (Boke and Mitchison, 2017; Bontems et al., 2009; Marlow and Mullins, 2008). The organization of the Bb is dynamically restructured as the oocyte matures: in early oocytes it is compacted, but upon maturation its contents disperse around the oocyte cortex in preparation for fertilisation (Bontems et al., 2009; Kar et al., 2025; Marlow and Mullins, 2008). Following fertilisation, these dispersed granules which are packed full of the proteins, RNAs and mitochondria necessary for the developing embryo, are reassembled at the cleavage furrows of the dividing blastomeres (Eno et al., 2019; Eno and Pelegri, 2018; Knaut et al., 2000). From here, this material is referred to as germ plasm. It is inherited asymmetrically and ultimately is found in four cells that will form the primordial germ cells (PGCs). This process of dispersal and reassembly is driven by liquid–liquid phase separation (LLPS) and aided by the reorganization of the microtubule cytoskeleton (Eno et al., 2019; Kar et al., 2025; So et al., 2021).

LLPS enables the formation of membraneless condensates that allow the segregation of proteins and RNAs from the cytoplasm without the need for a restricting lipid bilayer (Banani et al., 2017; Li et al., 2012). Within the germline, such condensates concentrate and protect germ plasm components, permit localised translation, and regulate RNA stability and processing (Hakes and Gavis, 2023; So et al., 2021; Updike and Strome, 2010). Their assembly depends on multivalent interactions provided by intrinsically disordered regions (IDRs), structured (RNA-binding) domains, and RNA, which can act as both a scaffold and a substrate for condensate formation (Banani et al., 2017; Li et al., 2012; Martin and Holehouse, 2020).

Integrity of germ plasm condensates and the Balbiani body depends on a core set of scaffold proteins. Chief amongst these in zebrafish is Bucky ball (Buc) (Bontems et al., 2009; Marlow and Mullins, 2008; Riemer et al., 2015). A functional homolog has also been described in *Xenopus laevis,* Xvelo (Boke et al., 2016; Nijjar and Woodland, 2013). Xvelo was shown to form amyloid-like aggregates that, unlike pathological amyloid-like aggregates formed by for instance FUS (Patel et al., 2015) or Tau (Wegmann et al., 2018), remain reversible and biologically active (Boke et al., 2016). A protein with analogous function to Buc and Xvelo in *Drosophila* is Oskar, which has been shown to be pivotal for germ plasm formation, and hence germ cell induction at the posterior pole of the *Drosophila* embryo (Ephrussi and Lehmann, 1992).

Buc aggregation is regulated by the Tudor-domain proteins Tdrd6a and Tdrd6b (Consorte et al., 2026; Roovers et al., 2018). Tdrd6a recognizes methylated arginines in Buc’s C- terminal region and regulates its mobility and prevents over-aggregation, thereby maintaining Bb and germ plasm dynamics (Roovers et al., 2018). Tdrd6b has been shown to interact with Buc via a Prion-like domain and, together with Tdrd6a is required for Buc condensation during early embryogenesis (Consorte et al., 2026). Indeed, the regulation and activity of Buc is a finely tuned process which involves myriad of other factors, such as the RNA-binding proteins Rbpms2 and Dazl, which recruit *buc* mRNA and other transcripts to the Bb (Heim et al., 2014; Krishnakumar et al., 2018), and microtubule-based cues that enable Buc condensate reorganization during oogenesis and egg activation (Campbell et al., 2015; Eno and Pelegri, 2018; Kar et al., 2025). Many aspects of germ plasm regulation remain unknown. Indeed, recent studies in other model systems such as *C. elegans* have identified many novel germ plasm factors that contribute to germ granule formation and behaviour (Chen et al., 2026; Hertz et al., 2022; Price et al., 2021), highlighting that key determinants are likely still left to be discovered. Of special interest for the work we describe here is *bucky ball-like* (*bucL*), a *buc* paralog present as a tandem duplicate in the non-teleost spotted gar which was further duplicated during the teleost genome duplication and then subject to widespread lineage-specific paralog loss (Škugor et al., 2016). Zebrafish retained *buc* alongside a single *bucL* paralog, *bucky ball 2-like* (*buc2l*). As Buc2l lacks the conserved N-terminal BUVE (Buc-Velo) motif shared by Buc and *Xenopus* Velo1, it was proposed not to function in germline formation (Škugor et al., 2016) and was not studied functionally. Our bioinformatic analyses reveal that Buc2l contains a tandem repeat motif whose copy number varies between species. Repeat-rich, disordered domains are a notable feature of proteins which mediate LLPS and condensate assembly, as each repeat can contribute an additional interaction site and thereby increase the overall valency of a protein (Banani et al., 2017; Li et al., 2012; Martin and Holehouse, 2020).

In this study, we investigated the biological role of Buc2l in zebrafish germ plasm organization and characterise its phase separation behaviour *in vitro*. We demonstrate that Buc2l forms assemblies whose growth depends on its tandem repeat region, interacts with known germ plasm components including Buc, and enhances Buc’s aggregation behaviour. We also show that loss of Buc2l *in vivo* prevents the formation of typical extensive Buc aggregates along the cleavage planes in the embryo, while still allowing the formation of small Buc granules. This is accompanied by loss of primordial germ cell specification. In absence of Buc2l the Bb was also affected, albeit less severely than germ plasm in the embryo, and it did not result in embryonic lethality as observed in embryos lacking Buc. Together, these findings identify Buc2l as a novel driver of higher-order germ plasm aggregation and establish a role for repeat-driven multivalency in this process.

## RESULTS

### Buc2l lacks a prion-like domain but contains a conserved tandem repeat region

To identify novel Buc interactors, we previously performed pull-down assays on zebrafish oocyte extracts followed by mass spectrometry (Roovers et al., 2018). Among the enriched proteins was a previously uncharacterized factor, Bucky ball 2-like (Buc2l). A schematic of the *buc2L* gene is displayed in Fig. 1A. Buc2l is a 962-amino-acid protein (Fig. 1B) with limited overall sequence similarity to Buc (18.2% identity, 28.4% similarity; EMBOSS Needle; Fig. S1A). Dot-plot analysis showed that this similarity was concentrated in the N-terminal ∼200 residues (Fig. S1B). IUPred predicted Buc2l to be largely intrinsically disordered, but unlike Buc it lacks a prion-like domain (PLD) (PLAAC, Fig. 1C). We raised a polyclonal antibody against the C-terminal region of Buc2l (amino acids 771–961). Immunoprecipitation from early zebrafish embryo extracts followed by mass spectrometry, using *buc2l* mutant (see below) extract as control, resulted in the expected enrichment of Buc2l and identified several known germ plasm components, including Buc, the PIWI protein Ziwi, and the Tudor-domain proteins Tdrd6a and Tdrd6b (Fig. 1D), confirming the previously identified Buc-Buc2l interaction.

**Figure 1.**
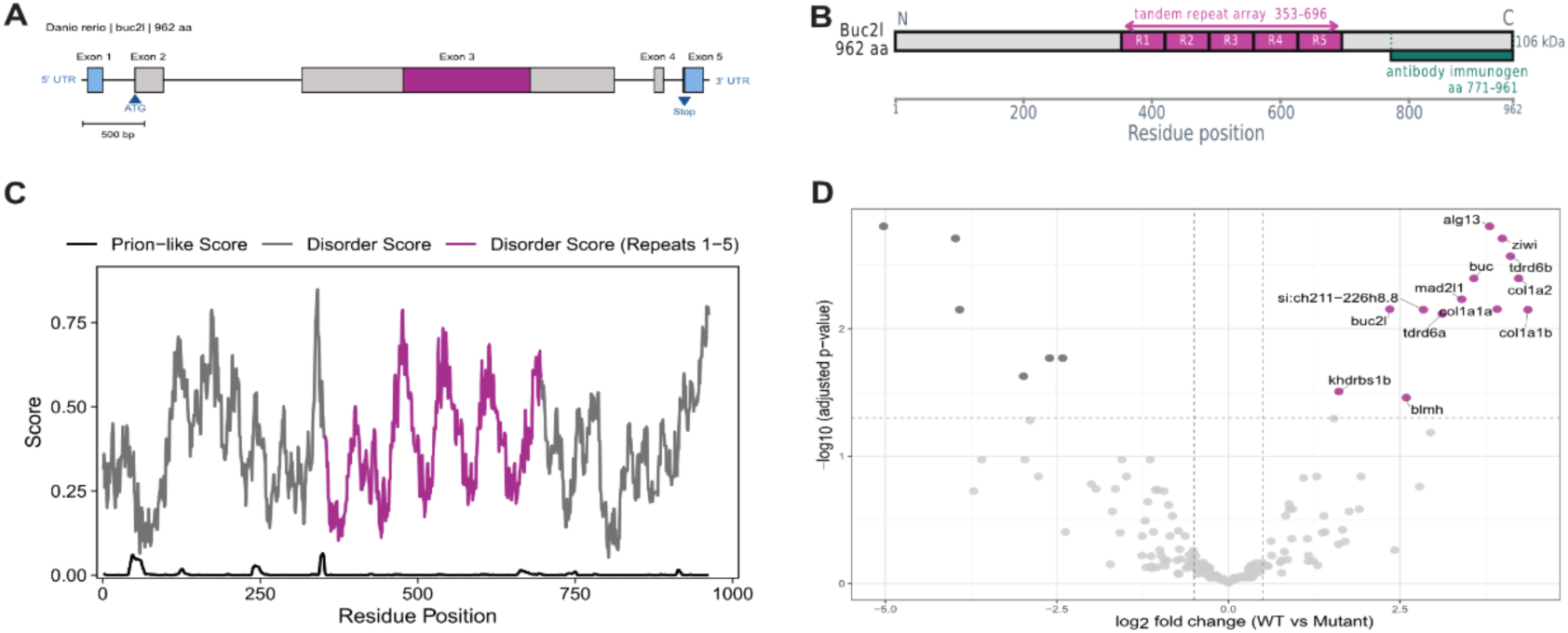
Buc2l is an intrinsically disordered, repeat-rich protein that associates with germplasm components. (A) Zebrafish buc2l gene structure. Scale bar: 500 bp. (B) Buc2l domain architecture. The tandem repeat array (residues 353-696) contains five ∼69 aa repeat units (R1-R5), and the region recognised by the anti-Buc2l antibody spans residues 771-961. (C) Predicted per-residue disorder (lUPred) and prion-like (PLAAC) propensity of Buc2l with the repeat region (R1-R5) highlighted. (D) Volcano plot of anti-Buc2l IP-MS from wild-type versus *buc2l* mutant embryos. Labeled proteins are significantly enriched in WT. Protein intensities were compared between genotypes with a linear model (limma) and Benjamini-Hochberg adjustment; n = 3 biological replicates per genotype. Proteins with logi fold change ≥ 1 and q < 0.05 were considered differentially enriched.

Sequence analysis of Buc2l protein revealed a block of ∼69 amino acids that repeated five times in tandem within the zebrafish Buc2l protein. We will refer to these blocks of amino acids as A-units. We compared Buc2l homologs across a range of teleost species and found that these A-units were conserved, being identified in multiple cyprinids (*Danio rerio*, *Pimephales promelas*, *Carassius auratus* and *Puntigrus tetrazona*), but also in the more basal *Anguilla anguilla* (Fig. 2A, B). Besides A-units, we also identified B-units, which are related in sequence to the A-units (Fig. S1C), but longer (∼167-residues per unit). These B-units were found in percids (*Perca fluviatilis*, *Sander lucioperca* and *Etheostoma cragini*) (Fig. 2A). The number of A- and B-units varied between species, from a single A- unit in the eel *A. anguilla* (Fig. 2B), to nine B-Units in perch (*P. fluviatilis*). The A- and B- units consistently started between residues ∼260 and 360, indicating that its position within the protein has been conserved even as repeat number and length varied (Fig. 2A). We note that some species have more than one *buc2l* copy (see below), explaining why for instance *Takifugu rubripes*, which is more closely related to percids than to cyprinids, appears twice in Fig. 2A, with one Buc2l protein carrying a single A-unit and another carrying a single B-unit.

**Figure 2.**
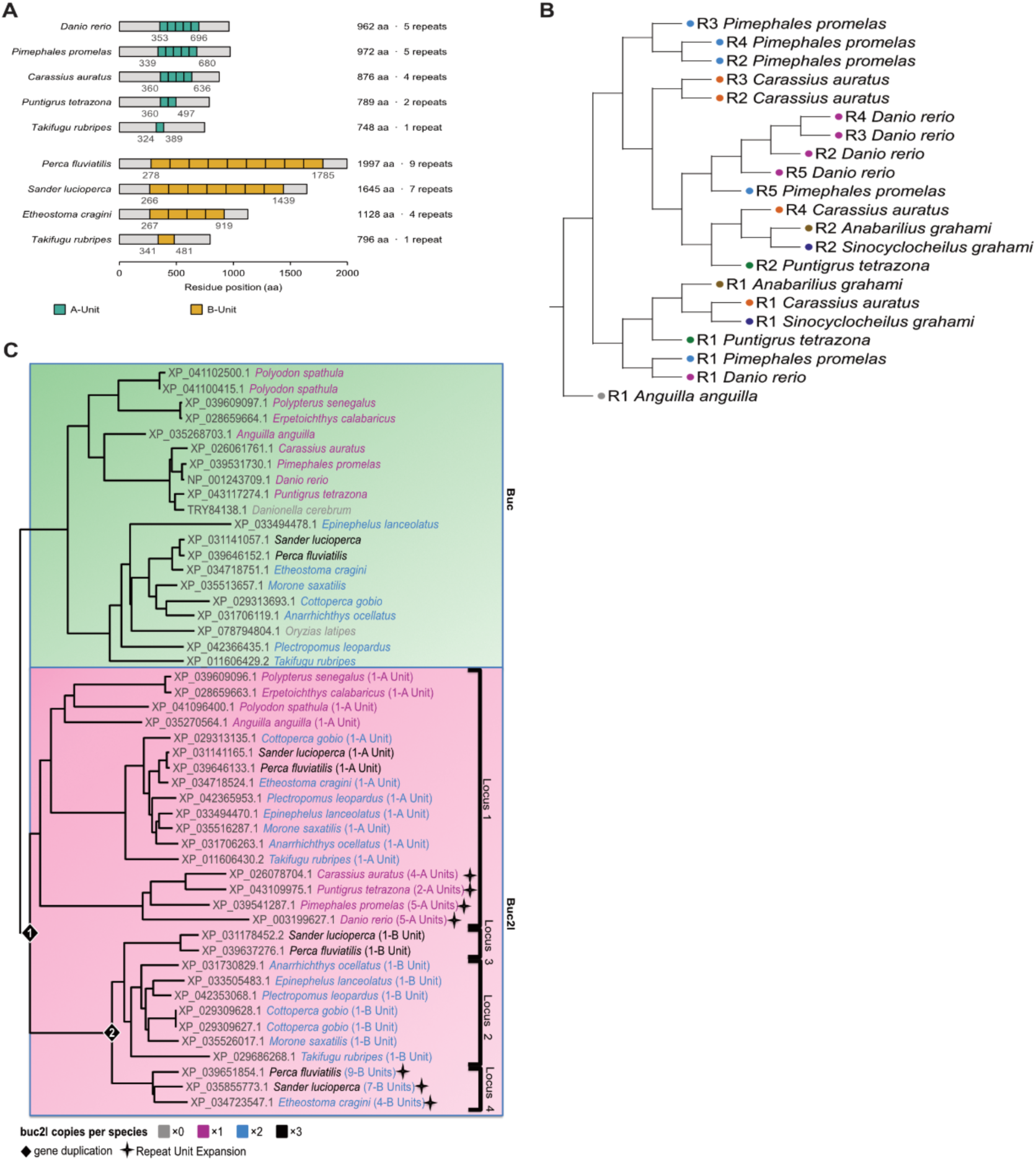
Buc2l repeat architecture and evolution in ray-finned fishes. (A) Domain architecture of Buc2l proteins carrying A-units (cyprinids) or B-units (percids). Both *T rubripes* Buc2l proteins are shown for comparison. Numbers mark the first and last residue of the repeat array. Protein length and repeat number are given at right. (B) Phylogeny of individual A-units (R1-R5), rooted on the single *A. anguilla* repeat. R1 units group across species, whereas R2-R5 group mainly by species, suggesting independent expansion of the repeat array in each lineage. (C) Phylogeny of Bug and Buc2l proteins. buc2l Loci 1-4 are defined by conserved flanking genes. Buc2l proteins with expanded A- or B-unit arrays are indicated; all other Buc2l proteins carry a single repeat unit.

To reconstruct the evolutionary history of the repeats, we built a phylogenetic tree of individual A-units, that are found in Buc2l in zebrafish, rooted on the single A-unit of *A. anguilla* (Fig. 2B). The presence of a single repeat in this early-diverging teleost suggests that ancestral Buc2l contained one repeat unit. Consistent with this, the first repeat unit (R1) of each species grouped together across species. In contrast, the remaining repeats grouped more by species rather than by position within the Buc2l protein (e*.g. D. rerio* repeat 3 grouped with other *D. rerio* repeats rather than with repeat 3 of *P. promelas*), consistent with lineage-specific duplication events. Thus, while repeat number expanded independently and to varying extents in different lineages, the position of the repeat region has been maintained.

To determine the relationship between *buc2l* genes carrying A- and B-units, we examined the genomic location of *buc2l* genes, assigning each to a locus based on its flanking genes (Fig. S2A). In early-diverging ray-finned fishes such as bichirs, paddlefish and eel, *buc* and *buc2l* lie side by side, suggesting that the two genes arose from a tandem duplication and were initially inherited together at the ancestral locus (locus 1; Fig. S2A, B). Although *buc* and *buc2l* were later separated in cyprinids, *buc2l* remained at locus 1 in all species examined that retain the gene (Fig. 2C; Fig. S2A). Notably, percomorphs acquired additional copies of *buc2l* through further gene duplications, and these copies are found at distinct genomic loci (loci 2–4). Locus 2 is found in several non-percid percomorphs, whereas loci 3 and 4 are restricted to percids (Fig. 2C; Fig. S2A). Of interest, the majority of Buc2l proteins carry only a single repeat unit, suggesting that this represents the ancestral state. At locus 1, this unit expanded in cyprinids (A-units), but not in percomorphs. Among the percomorph-specific paralogues, loci 2-4 are characterized by at least one B-unit. Loci 2 and 3 retained a single B-unit, whereas *buc2l* at locus 4 underwent an expansion of B-units (Fig. 2A; Fig. S1C). Thus, the Buc2l repeat region expanded at least twice independently, in different genes and in different lineages. Tandem repeats can increase the number of interaction sites, or valency, of a protein, and higher valency can promote the formation and further growth of biomolecular condensates (Banani et al., 2017; Li et al., 2012; Martin and Holehouse, 2020). The expansion of the Buc2l repeat units could therefore affect how strongly it may contribute to germ plasm assembly.

### Buc2l is a novel germ plasm-enriched protein that colocalises with Buc

Next we wanted to assess whether Buc2l, like Buc, is found in the Bb and/or the germ plasm. Using the antibody that was also used for IP-MS (see above), we performed immunostaining of zebrafish ovaries and found that Buc2l is enriched in the Balbiani body in both early and late-stage oocytes, exhibiting a spatial distribution closely resembling that of Buc (Fig. 3A). Buc2l was not enriched in the perinuclear nuage, where Tdrd6a, Tdrd6b and Ziwi are enriched (Consorte et al., 2026; Houwing et al., 2007; Roovers et al., 2018) (Fig. S3A). In early oocytes, Buc and Buc2l signals overlapped completely (Fig. 3A). In maturing oocytes, the characteristic dispersal of the Balbiani body was evident by Buc staining. Notably, Buc2l appeared to disperse earlier or more broadly than Buc, suggesting that a fraction of Buc2l may dissociate from the Balbiani body prior to the onset of full dispersal (Fig. 3B, Fig. S3B).

**Figure 3.**
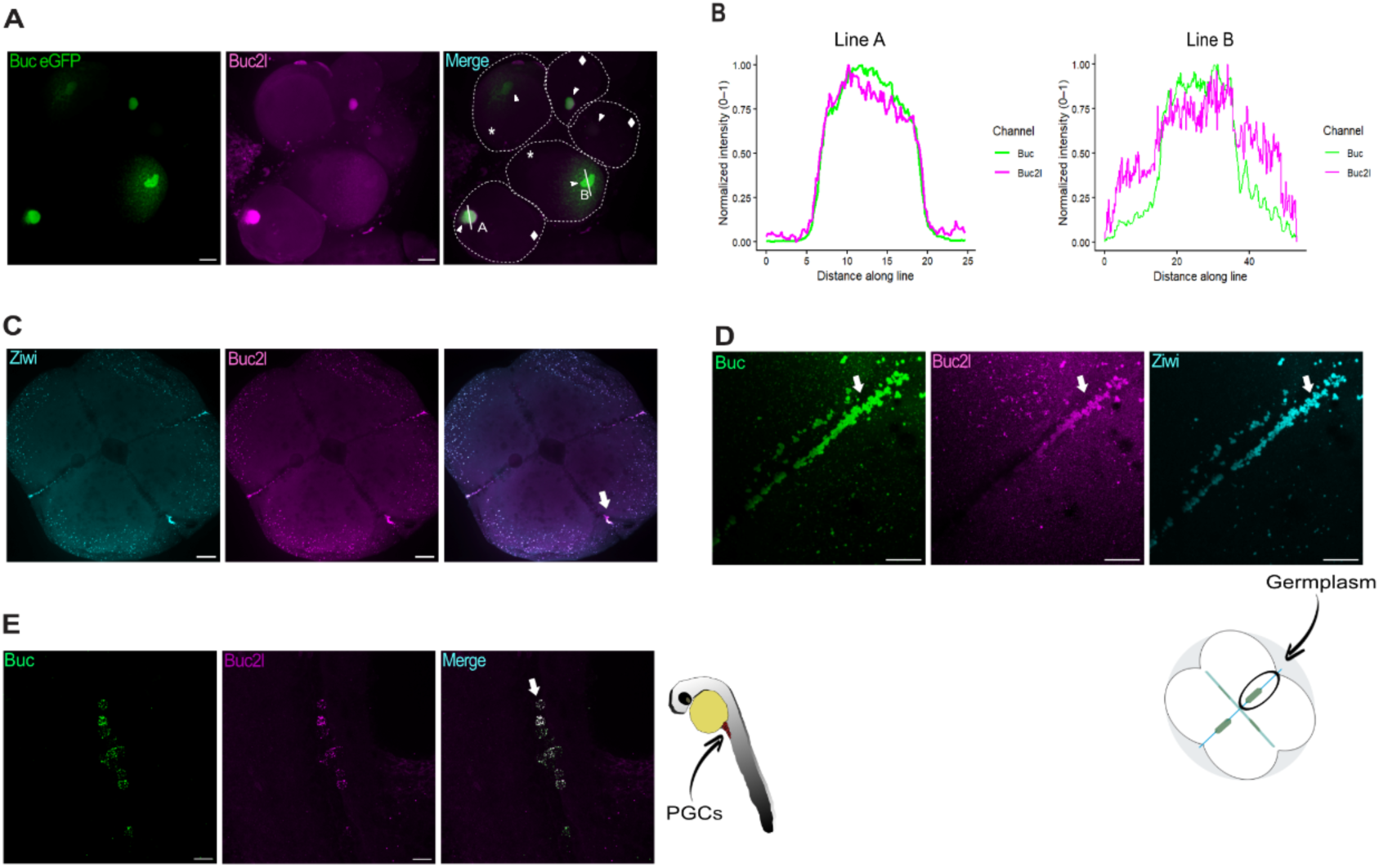
Buc2l colocalises with Buc in the Balbiani body, germplasm and primordial germ cells. (A) Oocytes from *Tg(buc.buc-eGFP)* ovaries immunostained for Buc2l. Oocytes are outlined; early (diamond) and late (asterisk) stage I oocytes, and the positions of the line profiles in B, are indicated. (B) Normalised intensity profiles of Buc-eGFP and Buc2l along line A (early stage I oocyte) and line B (late stage I oocyte) in A. (C) Overview of a 4-cell stage embryo immunostained for Ziwi and Buc2l. (D) Germplasm at the cleavage furrow of a 4-cell stage embryo immunostained for Buc, Buc2l and Ziwi; the schematic indicates the imaged region. (E) Primordial germ cells (PGCs) of a 24 hpf embryo immunostained for Buc and Buc2l; the schematic indicates the imaged region. The Balbiani body (A), germplasm (C, D) and PGCs (E) are indicated with pointers & arrows. Scale bars, 50 μm (C) and 20 μm (A, D, E).

Immunofluorescence analysis of 4-cell stage embryos revealed that Buc2l localises to numerous dispersed granules outside the furrow region (Fig. 3C) as well as to germ plasm at the cleavage furrows (Fig. 3C, D). Buc2l and Buc signals showed complete colocalization, and Buc2l additionally colocalised with other germline determinants such as Ziwi during this stage (Fig. 3C). At later stages (∼24 hpf), Buc2l puncta were detected within primordial germ cells (PGCs), demonstrating continued association with the germ cell lineage (Fig. 3E). Antibody specificity was validated by the absence of signal in oocytes (Fig. 4A) and embryos (Fig. S3C) derived from homozygous *buc2l* mutant females (see below).

**Figure 4.**
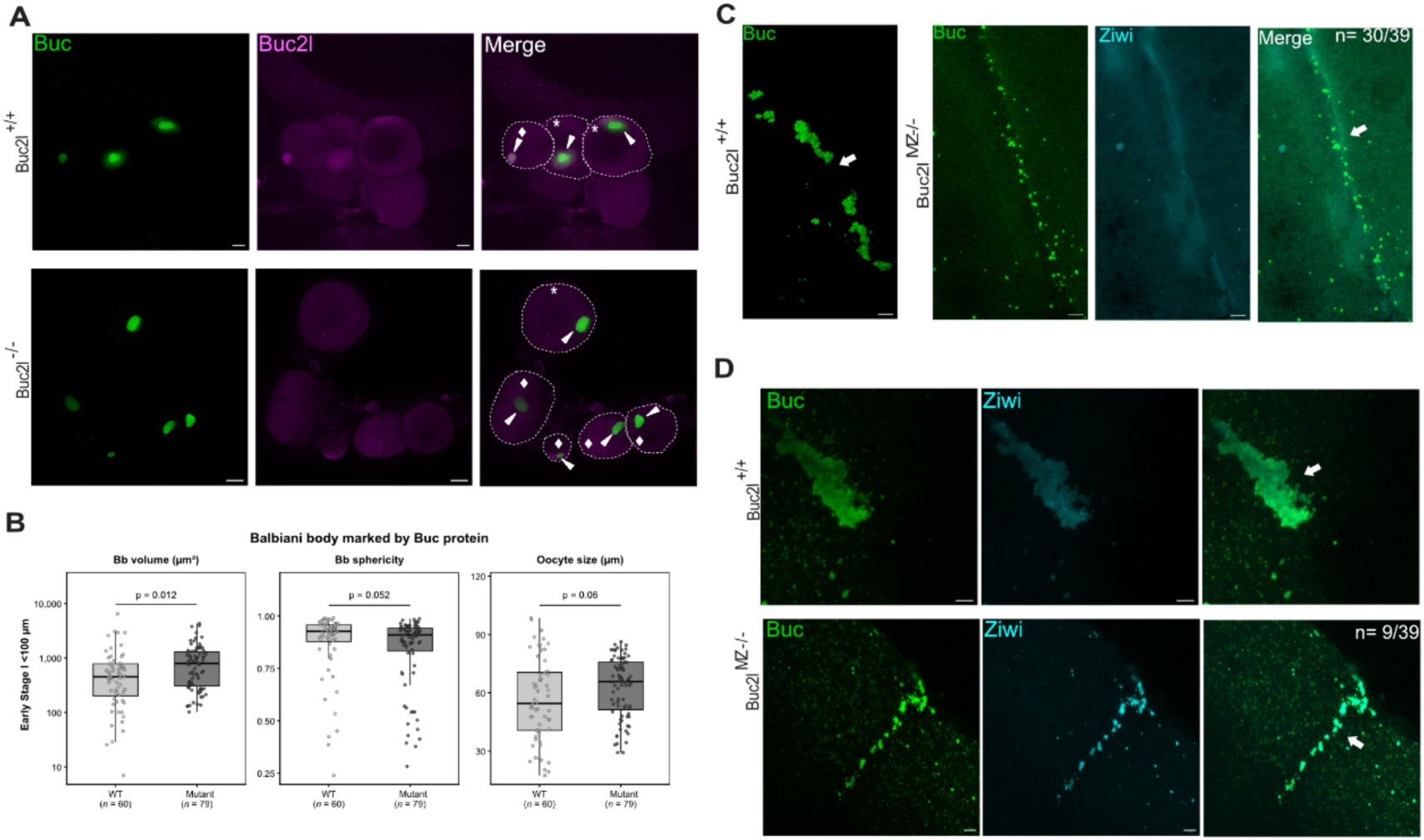
Loss of Buc2l alters the Balbiani body and impairs germplasm assembly in the embryo. (A) Oocytes from wild-type and *buc2l^-/-^* females immunostained for Buc and Buc2l, showing loss of Buc2l from the Balbiani body. Oocytes are outlined, and Balbiani bodies and oocyte stages are indicated. (B) Balbiani body volume, Balbiani body sphericity and oocyte size in early stage I oocytes (<100 μm) from wild-type (n = 60) and buc2l^-/-^ *(n* = 79) females, with the Balbiani body marked by Buc. Oocyte size is the mean of two perpendicular diameters.Each point is one oocyte; boxes show the median and interquartile range. Genotypes were compared by two-sided Mann-Whitney U test, Holm-corrected. (C) Buc in 4-cell stage wild-type and maternal-zygotic *buc2l^MZ-/-^* embryos, with Ziwi shown for the mutant n = 30/39 *buc2l^MZ-/-^* embryos exhibit no germplasm aggregation. (D) Buc and Ziwi at the cleavage furrow of 4-cell stage wild-type and *buc2^MZ-/-^* embryos, *n* = 9/39 *buc2l^MZ-/-^* embryos showed furrow-localised germplasm. Germplasm is indicated by arrows. Scale bars, 20 μm (A) and 5 μm (C, D).

### Buc2l is dispensable for oocyte development and Balbiani body formation

To investigate the function of Buc2l, we generated three CRISPR/Cas9-induced mutant alleles (N81, N101 and N140) (Fig. S4A). All three alleles disrupted the same splice site, producing similar aberrant transcripts (Fig. S4B) and internally deleted potential protein products (Fig. S4C). As all alleles exhibited comparable molecular and phenotypic characteristics, subsequent analyses were performed using the N101 allele.

Immunofluorescence confirmed loss of Buc2l protein from the Balbiani body in mutant oocytes (Fig. 4A). Balbiani body (Bb) morphology was compared between wild-type and mutant oocytes using volume and sphericity measurements (two-sided Mann-Whitney U test, Holm-corrected). As Bb morphology changes during oocyte growth, oocytes were binned by diameter into early stage I (< 100 µm, mature Bb) and late stage I (100–140 µm, Bb disassembly) (Elkouby and Mullins, 2017). In early stage I oocytes, Bbs marked by Buc were notably larger in *buc2l* mutants (median volume 794 versus 454 µm³, p = 0.012), while sphericity was not significantly different (median 0.91 versus 0.93; p = 0.052) (Fig. 4B). In contrast, when Bbs were marked by smFISH against *dazl* mRNA (Fig. S3D, E), early stage I Bbs were comparable in volume between genotypes (median 895 versus 1,177 µm³; p = 0.14), although mutant Bbs were markedly more spherical (median 0.93 versus 0.77; p = 0.008); mutant oocytes in this group were also smaller (median 70 versus 81 µm; p = 0.013). Intriguingly, the most striking difference emerged during Bb disassembly. In late stage I oocytes of comparable size (median 105 versus 111 µm; p = 0.10), *dazl*-marked Bbs were smaller in *buc2l* mutants (median volume 2,770 versus 9,302 µm³; p = 0.003) and remained more compact (median sphericity 0.56 versus 0.39; p = 0.045), whereas wild-type Bbs expanded and became markedly irregular (Fig. S3D, E). These contrasting phenotypes between the Bb when assessed via Buc protein versus *dazl* mRNA suggest that loss of Buc2l causes an uncoupling of the protein and RNA components of the Balbiani body, implying a role for Buc2l in coordinating RNA integration into the Bb and/or a role for mRNA in Bb expansion during disassembly.

### Buc2l is required for germ plasm assembly in the embryo

Striking defects in germ plasm architecture were observed in embryos obtained from homozygous mutant mothers (*buc2l^MZ-/-^*). In wild-type embryos at the 4-cell stage, Buc- containing granules accumulate at the cleavage furrows and coalesce into large germ plasm structures (Fig. 4C). In *buc2l^MZ-/-^* mutants, Buc was still recruited to cleavage furrows, but the granules failed to coalesce into large germ plasm structures (Fig. 4C). Instead, they appeared as small, scattered granules along the furrow. Consistent with this, Ziwi, which normally accumulates in germ plasm together with Buc, was largely diffuse throughout the cytoplasm of mutant embryos, with only weak enrichment along the furrow (Fig. 4C). Interestingly, while most imaged embryos displayed a widespread loss of germ plasm foci (30/39), a small number retained granules at certain furrows (9/39 embryos; Fig. 4D). In these embryos, the germ plasm markers still overlapped with Buc, indicating that the residual granules represent genuine, if reduced, germ plasm. Also, the Tdrd6a and Tdrd6b proteins lost their recruitment to germ plasm structures in absence of Buc2L (Fig. S3C). In addition, *dazl* and *buc* mRNAs were not detectably enriched at the cleavage furrows of *buc2l*^MZ−/−^ embryos, even when traces of Buc granules were detectable (Fig. S3F). Together, these data show that Buc2l is dispensable for the formation of individual Buc-containing granules and their recruitment to the cleavage furrows, but is essential for coalescing them into large germ plasm structures. Additionally, Buc2L promotes the recruitment of other germ plasm components, such as Ziwi, Tdrd6a and Tdrd6b and germ plasm mRNAs.

### Loss of germ plasm impairs primordial germ cell development

We next examined primordial germ cell (PGC) development in *buc2l^MZ-/-^*mutants. At 24 hpf, *buc2l^MZ-/-^* embryos exhibited a near complete loss of PGCs, as assessed using the Tg(vasa:eGFP) transgenic line (Fig. 5A, B). The few germ cells that were detected were frequently ectopically positioned and failed to migrate to the gonadal ridge, instead remaining scattered throughout the embryo (Fig. 5C). Despite these germline defects, overall embryonic morphology appeared largely normal, indicating that the loss of Buc2l did not broadly disrupt embryonic development. When raised to adulthood, *buc2l^MZ-/-^*mutants developed exclusively into sterile males and lacked clear testes (Fig. 5D, E). This phenotype is consistent with the previously established link between PGC loss and male- only sex determination in zebrafish, whereby the lack of a sufficient germ cell population during the critical window of sex determination causes the fish to develop as a male (Siegfried and Nüsslein-Volhard, 2008; Slanchev et al., 2005)

**Figure 5.**
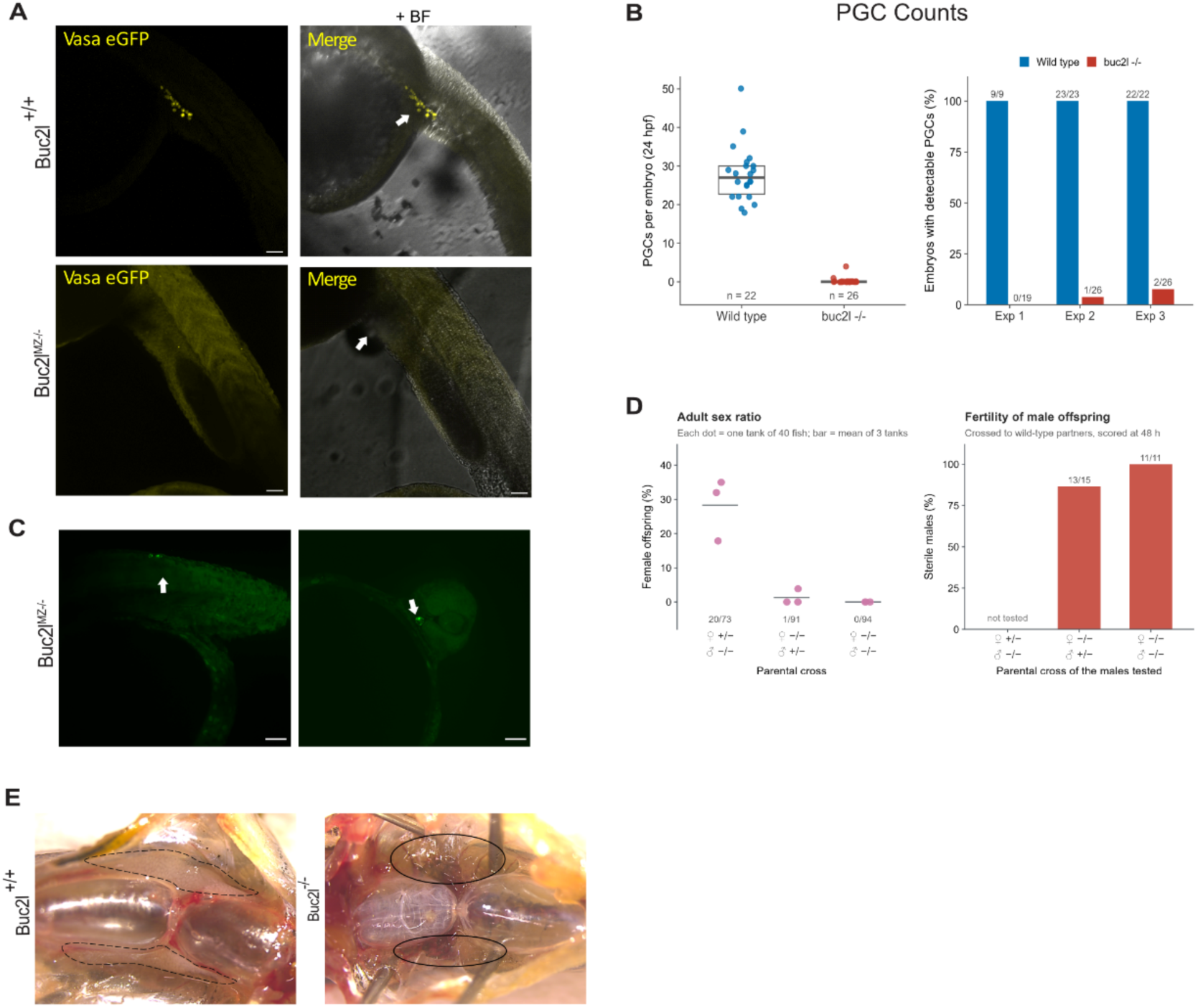
Loss of Buc2l results in loss of primordial germ cells and sterile male offspring. (A) *Tg(vasa:eGFP)* embryos at 24 hpf from wild-type and *buc2l^-/-^* females, showing Vasa-eGFP alone (left) and merged with brightfield (right). PGCs are indicated. (B) PGCs per embryo at 24 hpf in wild-type (n = 22) and *buc2l^MZ-/-^* (n = 26) embryos from one experiment (left). Boxes show the median and interquartile range. Percentage of embryos with detectable PGCs in three independent experiments (right), numbers give embryos with PGCs / total embryos. (C) Mislocalised PGCs in *buc2l^M-/-^* embryos at 24 hpf, in the trunk muscle (left) and head (right). (D) Percentage of female offspring from the indicated parental crosses (left). Each point is one tank of 40 fish, and lines show the mean of three tanks. Percentage of sterile males among offspring from the indicated parental crosses (right), assessed by crossing to wild-type partners and scoring fertilisation at 48 h. Numbers give females / total fish (left) and sterile males / males tested (right). (E) Dissected adult wild-type and *buc2F* males, with gonads indicated. Scale bars 20 μm (A, C).

Taken together, these results establish a role for Buc2l in germ plasm biology. Buc2l is not required for Balbiani body formation per se as Balbiani bodies still assemble in its absence, even if their size and content may be affected. Buc2l also appears dispensable for the initial recruitment of germ plasm components to cleavage furrows but is essential for the higher-order condensation of these components into structurally organised and functional germ plasm during early embryogenesis. In the absence of this organization PGC specification is severely compromised.

### Buc2l and Buc co-assemble into large assemblies in vitro

To investigate the physical properties of Buc2l and its interaction with Buc, we expressed and purified recombinant proteins under denaturing conditions, including ectopic fluorescent labelling, (Fig. S6A, B) and diluted them into non-denaturing buffer to 500 nM, to assess their condensation behaviour. Each protein alone formed assemblies (Fig. 6A, B) and assembly was concentration-dependent, with both proteins forming markedly larger assemblies at ≥1000 nM, especially Buc2l (Fig. S5A).

**Figure 6.**
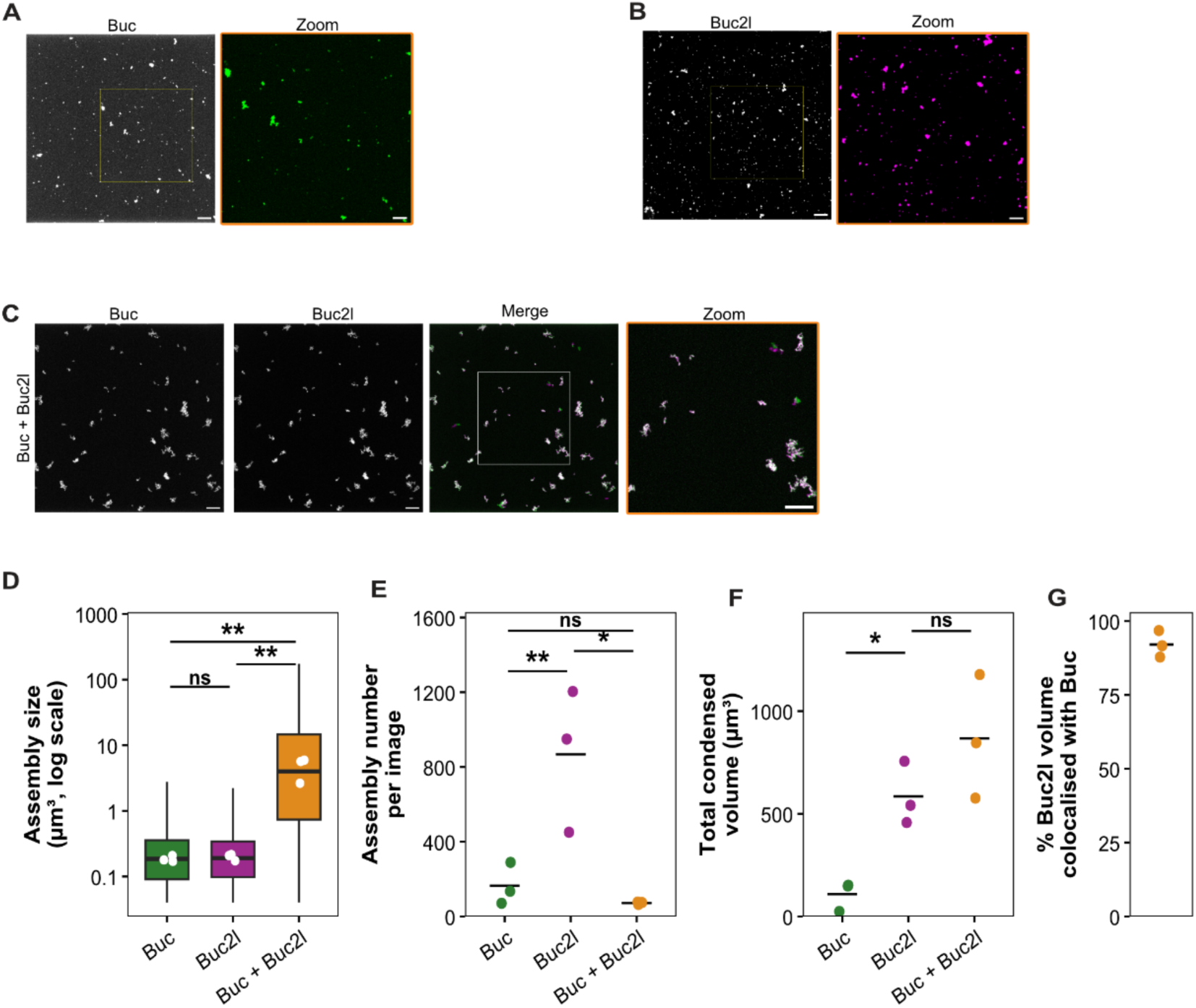
Buc and Buc2l co-assemble into large assemblies in vitro. (A, B) Representative images of recombinant Buc (A) and Buc2l (B), each at 500 nM; boxed regions are enlarged on the right. (C) Buc and Buc2l combined (500 nM each): Buc and Buc2l channels, merge and zoom of the boxed region. Images are maximum-intensity projections of confocal z-stacks. (D- G) Quantification of assemblies formed by Buc alone, Buc2l alone and Buc + Buc2l; for the mixture, assemblies were segmented in the Buc2l channel. (D) Assembly size (volume, log scale). Boxes show the median and interquartile range of all assemblies; whiskers, 1.5× IQR; white dots, replicate medians. (E) Assemblies per field. (F) Total assembly volume per field. (G) Percentage of Buc2l assembly volume overlapping Buc assemblies in the mixture. For E-G, values were calculated per field and averaged within each replicate (6-13 fields per condition per experiment); dots, replicate means; lines, mean of replicates; n - 3 independent experiments. Statistics were performed on replicate values (medians for D, means for E-G; log scale for D-F). Buc vs Buc2l: linear model with experiment as block, Holm-corrected. Alone vs mixture: Welch’s t-test. Scale bars, 10 μm; zooms, 5 μm. *p < 0.05, **p < 0.01; ns, not significant.

At 500 nM, Buc and Buc2l assemblies were of comparable size, but Buc2l formed roughly five times more of them, resulting in a higher total condensed volume for Buc2l (Fig. 6D– F). When Buc and Buc2l were combined (500 nM each), they formed larger assemblies than either Buc or Buc2l alone, while the number of assemblies decreased (Fig. 6C,D). This increase in size was observed in both the Buc and the Buc2l fluorescence signals, and most of the Buc2l signal overlapped with Buc (∼93%, Fig. 6G), indicating that the two proteins assembled into shared structures rather than forming separate populations. Together, these data show that Buc and Buc2l cooperate to build large assemblies that neither protein forms efficiently on its own, mirroring the observation of only small Buc aggregates in *buc2l* mutants *in vivo* (Fig. 4C, D).

### Assembly size scales with Buc2l A-unit number

To test the contribution of the Buc2l A-units to its aggregation behaviour, we investigated a series of Buc2l variants with varying numbers of A-units. Specifically, we deleted unit 5 (Δ5), units 3-5 (Δ3-5), units 2-5 (Δ2-5) and a variant lacking all five A-units (Δ1-5). On their own, all variants still formed assemblies, albeit fewer and smaller than their full-length counterpart, leading to lower overall condensed volume (Fig. 7A-D). Interestingly, the mean assembly size scaled with the number of A-units retained: more units led to larger assemblies (Fig. 7B). We also found that at ≥1000 nM wild-type Buc2l started to form very large aggregates, and that Buc2l(Δ1-5) failed to do so (Fig. S5A). Thus, we conclude that the A-units of Buc2l control Buc2l aggregate size.

**Figure 7.**
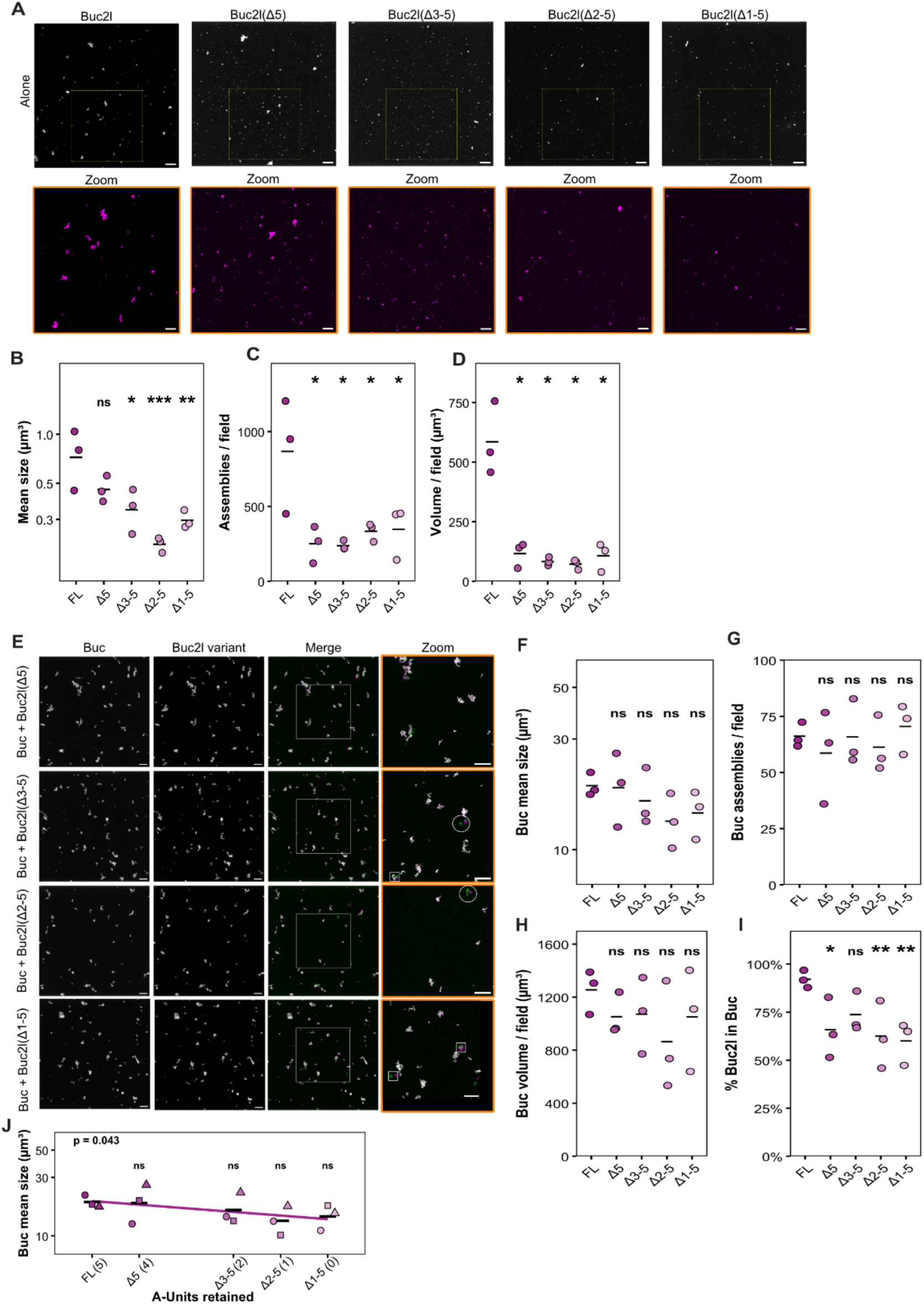
Buc2l A-units drive assembly growth. (A) Buc2l (FL) and A-unit deletions (500 nM) alone; zooms below. (B-D) Mean assembly volume (B), assemblies per field (C), total volume per field (D). (E) Buc with each Buc2l variant (500 nM each). In the zooms, circles mark Buc2l assemblies adjacent to Buc without merging; squares mark objects offset by drift during acquisition, excluded from quantification. (F-1) Buc mean assembly volume (F), Buc assemblies per field (G), Buc volume per field (H), % Buc2l-variant volume within Buc/Buc2l assemblies (I). (J) The data in (F) replotted against A-units retained, with fitted trend; t-test on slope, uncorrected. Scale bars, 10 μm; zooms, 5 μm. Shapes, replicate means; lines, mean; *n* = 3. Linear model with replicate as block on log™ values, Holm- corrected; *p < 0.05, “p < 0.01, ***p < 0.001 vs FL; ns, not significant.

We next combined each variant with Buc (500 nM each; Fig. 7E). None of the individual variants differed significantly from full-length Buc2l in the size (0.70–0.98×), number (0.85–1.06×) or total volume (0.65–0.84×) of the resulting Buc assemblies (Fig. 7F-H). However, the data did reveal a significant linear trend (p = 0.043; Fig. 7J) that was consistent with the effect of A-unit deletion on the aggregation of Buc2l alone, suggesting that the effect of the A-units is dampened, but still significant when Buc and Buc2l form co-assemblies. Using the overlapped volume ratio between segmented assemblies, we observed that all deletion variants were efficiently incorporated into Buc assemblies, albeit less efficiently than full length Buc2l (Buc2l (Δ1-5) ∼60%, versus ∼92% of Buc2l colocalized with Buc, Fig. 7I). Objects in which the two channels were offset by drift during acquisition were excluded from this analysis.

We also tested the A-unit region of Buc2l alone, separated from its normal N- and C- terminal tails (Buc2l (R1-5)). Interestingly, this protein only formed small aggregates. When mixed with Buc, it was able to accumulate in Buc granules, but it did not stimulate their growth like Buc2l does (Fig. S7C, D). We conclude that the Buc2l A-units drive assembly size of Buc-Buc2l aggregates, but only in context of full length Buc2l protein. The A-units also contribute to, but are not required for the interaction between Buc and Buc2l. Likewise, the protein that is potentially encoded by the *buc2l(N101)* allele (Δ78– 873; Fig. S4C, S6) was efficiently recruited into Buc assemblies (∼95%), but these assemblies were ∼5x smaller and more numerous than those formed with full-length Buc2l (Fig. S7C, D), mirroring the small, scattered Buc granules seen in *buc2l^MZ−/−^* embryos (Fig. 4C).

### A Buc-Buc2l interaction that can drive condensation

Using AlphaFold3 (Abramson et al., 2024) to model Buc–Buc2l interactions, we found a potential weak interaction between a C-terminal alpha-helix in Buc2l (residues ∼890-962) and a short N-terminal alpha-helix in Buc (residues ∼125-140). Even though the predicted interaction was only weak and would normally not be considered significant (ipTM=0.29; pTM=0.42) we probed this experimentally. We generated recombinant proteins covering the C-terminal segment of Buc2l (697-962) and an N-terminal fragment of Buc (121-325) (Fig. S6). Neither fragment alone formed assemblies on its own (Fig. S7A, B, E). However, when combined (500 nM each) they formed assemblies containing both proteins (Fig. S7F), suggesting that these regions can indeed interact, albeit weakly. Consistent with a role for the Buc2l C-terminus in this interaction, the C-terminal fragment was efficiently recruited into assemblies formed with full-length Buc (∼92% of its volume), although the resulting Buc assemblies were smaller and more numerous than those formed with full- length Buc2l (Fig. S7C, D).

## DISCUSSION

In this study, we identify Buc2l as a novel component of the zebrafish germ plasm, playing a critical role in germ plasm assembly during early embryogenesis. *In vitro*, Buc2l forms assemblies on its own, and together with Buc it forms assemblies far larger than either protein forms alone. *In vivo*, Buc2l is essential for the aggregation of Buc, Ziwi, Tdrd6a/b and germ plasm mRNAs into the large germ plasm structures that form along the cleavage furrows of wild-type embryos. In its absence, germ plasm remains fragmented, primordial germ cells are almost entirely lost, and the resulting fish develop as sterile males. Notably, Buc2l is not required for Balbiani body formation or for oogenesis, although its loss does affect Bb organization. Its potential functions in both the oocyte and the embryo will next be discussed, followed by a discussion on the potential implications of our results on the A-unit repeats that are present within the Buc2l protein.

Buc2l is strongly enriched in the Balbiani body from early oocyte stages and colocalizes with Buc. Yet, its loss does not prevent Bb formation, oogenesis or female fertility. The Bb is nevertheless affected. In *buc2l* mutants, the Buc-marked Bb is enlarged in early stage I oocytes, whereas the dazl-marked Bb is more compact at this stage and both smaller and more compact during Bb disassembly in late stage I oocytes. This suggests that Buc2l is not essential for forming the Bb but does contribute to a compact Bb in which protein and RNA components are properly integrated. One possible explanation for why the Bb still forms in absence of Buc2l is that, within the nuclear cleft where the Bb first develops, the local concentration of Buc is high enough to drive assembly without the additional valency provided by Buc2l. In the embryo, by contrast, germ plasm must be rapidly reassembled from dispersed cortical granules during cell division and cytoplasmic streaming, a situation in which efficient coalescence of small Buc granules may be much more challenging and thus possibly more dependent on enhancers of aggregation, like Buc2l. Intriguingly, during oocyte maturation, Buc2l appears to dissociate from the Bb earlier than Buc, raising the possibility that Bb dispersal may be made more effective by the loss of Buc2l, in line with the molecular role we demonstrate for Buc2l: it enhances Buc aggregation. This may also explain why the Bb, as judged by Buc protein, is slightly larger in absence of Buc2l. In contrast, the *dazl*-marked Bb failed to expand in *buc2l* mutants during disassembly. This may reflect a defect in the recruitment of mRNPs to the Bb, or in their release during Bb dispersal. Whether Buc2l is directly responsible for this defect, or whether this represents an indirect effect of changes in Buc aggregation state is currently unclear. This also applies to the defects in Ziwi, Tdrd6a/b and mRNP recruitment in the embryo. The primary effect of Buc2l loss in embryos may again be on Buc granule aggregation, followed by a loss of other components that may require a certain critical mass of Buc aggregation. However, we can also not exclude a direct role for Buc2l in the recruitment of these additional factors to the germ plasm.

The failure of germ plasm assembly in *buc2l*^MZ−/−^ embryos had severe consequences for the germline. PGCs were almost entirely absent at 24 hpf, and the few that formed were frequently mispositioned. This is consistent with the critical requirement for germ plasm in PGC specification (Bontems et al., 2009; Consorte et al., 2026) and suggests that fragmented germ plasm, even when its components reach the cleavage furrows, cannot support germ cell fate.

A striking feature of Buc2l is its tandem repeat region formed by five distinct A-units. Our deletion analysis suggests that these repeated units provide a source of multivalency: the more repeats were present the larger the Buc-Buc2l assembly grew. The isolated repeat region formed small condensates on its own, but did not support the growth of Buc condensates, suggesting that this property depends on some synergy with other regions of the Buc2l protein. Indeed, Buc2l lacking all five repeats also condensed readily, indicating that the ability to condense is distributed across the protein rather than confined to the repeats. The repeats did, however, contribute to the interaction with Buc itself: variants lacking A-units were recruited into Buc assemblies less efficiently than full-length Buc2l, without being excluded from them. How the A-unit repeats mediate growth of Buc condensates currently remains unclear.

The repeat region of Buc2l has expanded independently in different teleost lineages, while its position within the protein has been maintained. The single repeat in *A. anguilla* and the grouping of the first repeat across species suggest that ancestral Buc2l contained a single repeat, and that additional copies arose through lineage-specific duplications. Synteny analysis further indicates that *buc* and *buc2l* arose from a tandem duplication at an ancestral locus, and that the repeat region expanded independently at least twice: in cyprinids at the ancestral locus, giving rise to the A-unit repeats, and in percids in a paralogue at another locus, giving rise to the B-unit repeats. The conservation of repeat sequence alongside variable repeat number suggests that selection acts on the structural unit itself rather than on a specific valency. Of particular interest, an analogous architecture has recently been described for PINT-1, a disordered germ granule protein in *C. elegans* that combines a conserved, structured N-terminus with a C-terminal intrinsically disordered region containing four tandem repeats, the number of which differs across nematode species (Chen et al., 2026). In both proteins, a conserved N-terminal region is paired with a variable, repeat-rich C-terminus, suggesting that selection preserves the core of the protein (which holds the ancestral function of the family) while permitting variation in the repeat domains (which modify the ancestral function). As repeat number influenced assembly size *in vitro*, changes in copy number could provide a means to tune germ plasm assembly between species to adapt to varying conditions. Comparing Buc2l orthologues with different repeat numbers would be a promising way to test this directly.

### Author contributions

R.F.K. and F.C. designed the study. F.C. performed the immunostaining, smFISH, microscopy, in vitro assembly assays and image analysis. M.M.M expressed, purified and labelled the recombinant proteins. M.A. carried out the repeat annotation and phylogenetic analyses. W.B. designed and generated the CRISPR alleles and Y.E.S. performed the injections. F.C. wrote the manuscript with input from R.F.K.

## Acknowledgments

We thank the Microscopy and Histology Core Facility of the Institute of Molecular Biology (IMB), Mainz, for support with imaging; the Protein Production Core Facility for recombinant protein expression, purification and labelling; and the Proteomics Core Facility for mass spectrometry and data analysis. We thank all members of the Ketting lab for helpful discussions, with special thanks to Nadine Wittkopp and Joanna Michowicz, and the IMB Fish Facility for their excellent zebrafish care.

## Data availability

The mass spectrometry raw data have been deposited to the MassIVE data repository with the dataset identifier PXD087500. Reviewer Access: https://proteomecentral.proteomexchange.org/ui?pxid=PXD087500 ftp://MSV000104040@massive-ftp.ucsd.edu

## MATERIALS AND METHODS

### Computational sequence analyses

Homologous protein sequences were collected from relevant species using sequence similarity searches using NCBI BLASTp (Sayers et al., 2025). Multiple sequence alignments of full sequences or repeats were constructed using MUSCLE (Edgar, 2004) via the EBI web alignment service (Madeira et al., 2024). Phylogenetic trees were produced with ClustalX (Larkin et al., 2007) and represented with NJplot (Perrière and Gouy, 1996).

### Zebrafish husbandry

Danio rerio were maintained under standard laboratory conditions at 28°C on a 14 h light / 10 h dark cycle. Fish were cared for in accordance with the German Animal Welfare Act. All experimental procedures were performed in compliance with institutional and governmental regulations.

Embryos were obtained by natural mating and collected in E3 embryo medium (5 mM NaCl, 0.17 mM KCl, 0.33 mM CaCl₂, 0.33 mM MgSO₄). Embryos and larvae were maintained at 28.5°C in a temperature-controlled incubator up to 5 days post fertilisation.

All lines and alleles used in this study are listed in Table 2. *buc2l* mutant lines were generated in wild-type AB or Tübingen (TU) and periodically outcrossed to an AB/TU background and maintained as heterozygous carriers. Homozygous *buc2l*^-/-^adults were obtained by incrossing heterozygous carriers.

### Generation of buc2l mutant alleles

One-cell stage wild-type (AB/TU) embryos were injected with ∼1 nL of a mixture containing Cas9 mRNA and two sgRNAs. Injected embryos were raised at 28°C to adulthood and outcrossed to wild-type fish, and F1 offspring were screened for deletions at the target locus by PCR (see Genotyping) and Sanger sequencing. Three alleles were recovered: N81 (907 bp deletion), N101 (399 bp deletion) and N140 (748 bp deletion with a 1 bp insertion). All three remove the intron 2–exon 3 boundary, resulting in aberrant splicing and an in-frame deletion of residues 78–873.

### Genotyping

Genomic DNA was extracted from the tail fin of anaesthetised adult zebrafish. Fin clips were incubated in lysis buffer (50 mM KCl, 2.5 mM MgCl₂, 10 mM Tris-HCl pH 8.0, 0.5% NP-40, 0.5% Tween-20, 0.01% w/v gelatin, 0.1 mg/mL Proteinase K) for 1 hour at 60°C, followed by inactivation at 95°C for 10 minutes. The lysate was used directly as a PCR template.

*Buc2l* mutant alleles were genotyped using primers spanning the intron 1- exon 2 boundary. Primer WB1271 (5’-GGG TCG TGG TTA AGC CCA AAG ACA TG-3’) binds within intron 1, while primers WB1272 (5’-CTT AAC ATC CCA GCT TCA ACC CTC TG-3’) and WB1267 (5’-CAT GTG GCT CCA GCT ACT GC-3’) both bind within exon 2. PCR products were analysed by agarose gel electrophoresis (2% agarose, 1× TAE buffer).

### Generation and purification of anti-Buc2l antibodies

Two polyclonal antibodies against Danio rerio Buc2l were raised in rabbits (Eurogentec, Belgium) using the “speedy protocol” immunisation schedule. Rabbits were immunised with recombinant His-MBP-3C-tagged Buc2l C-terminus (amino acids 771–961).

For affinity purification, GST–Buc2l-C (771–961) antigen was coupled to SulfoLink coupling resin via free cysteine residues in the presence of 2 mM TCEP. Immune sera were filtered and applied to the antigen-coupled columns overnight at 4°C. Columns were washed with PBS and PBS + 0.1% Triton X-100, and antibodies were eluted with 150 mM NaCl, 200 mM glycine pH 2.3. Eluates were neutralised, rebuffered into PBS + 10% glycerol + 0.05% NaN₃, concentrated to 1 mg/mL, aliquoted, and stored at −80°C.

### Immunoprecipitation and mass spectrometry

Immunoprecipitation coupled to mass spectrometry (IP–MS) was performed using zebrafish embryos at the 4-cell stage. Embryos were collected immediately after fertilisation, staged accordingly, and dechorionated using Pronase. Approximately 50 embryos per condition (wild type vs. buc2l N101) were washed in phosphate-buffered saline (PBS) and centrifuged at 500 × g for 5 min at 4°C. Embryos were resuspended in ∼500 μL lysis buffer (25 mM Tris-HCl pH 7.5, 150 mM NaCl, 1.5 mM MgCl₂, 1% Triton X- 100, protease inhibitor cocktail, and 1 mM DTT freshly added) and lysed on ice.

Protein G magnetic beads (30 μL slurry per immunoprecipitation) were washed three times with 1 mL lysis buffer by gentle rotation at 4°C for 2 min per wash, followed by magnetic separation. Beads were then incubated with primary antibody at a ratio of 1 μg antibody per 10 μL beads in 500 μL lysis buffer. Clarified embryo lysates were added to the antibody-coupled beads, and 1% of total lysate volume was retained as input control. Samples were incubated overnight at 4°C with rotation. Following incubation, the unbound fraction was collected, and beads were washed five times for 5 min each with wash buffer (25 mM Tris-HCl pH 7.5, 300 mM NaCl, 1.5 mM MgCl₂, supplemented with freshly added DTT), with magnetic separation between washes.

Bound proteins were eluted by resuspending beads in 18 μL NuPAGE LDS sample buffer (1×, without DTT) and incubating at 70°C for 10 min. Eluates were collected after brief centrifugation.

### Enzymatic protein digestion

Samples were processed using the SP3 approach (Hughes et al., 2019). Proteins were reduced in 5 mM DTT, alkylated in 15 mM iodoacetamide in the dark and quenched in 5 mM DTT. Enzymatic protein digestion was performed using trypsin at 37°C overnight. Following acidification by formic acid, the peptides were purified by solid phase extraction in a C18 StageTip format (Rappsilber et al., 2003)

### Liquid chromatography tandem mass spectrometry

Peptides were separated via an in-house packed 45-cm analytical column (inner diameter: 75 μm; ReproSil-Pur 120 C18-AQ 1.9-μm silica particles, Dr. Maisch GmbH) on a Vanquish Neo UHPLC system (Thermo Fisher Scientific). Online reverse phase chromatography was performed through a 70-min gradient of 1.6-32% acetonitrile with 0.1% formic acid at a nanoflow rate of 300 nl/min. The eluted peptides were sprayed directly by electrospray ionization into an Orbitrap Astral mass spectrometer (Thermo Fisher Scientific). Mass spectrometry was conducted in data-dependent acquisition (DDA) mode using a top50 method with one full scan in the Orbitrap analyser (scan range: 325 to 1,300 m/z; resolution: 120,000, target value: 3 × 106, maximum injection time: 25 ms) followed by 50 fragment scans in the Astral analyser via higher energy collision dissociation (HCD; normalized collision energy: 26%, scan range: 150 to 2,000 m/z, target value: 1 × 104, maximum injection time: 10 ms, isolation window: 1.4 m/z). Precursor ions of unassigned, +1 or higher than +6 charge state were rejected. Additionally, precursor ions already isolated for fragmentation were dynamically excluded for 15 s.

### Mass spectrometry data processing and statistical analysis

Mass spectrometry raw data were processed in MaxQuant software (version 2.6.6.0) (Cox and Mann, 2008) using its built-in Andromeda search engine (Cox et al., 2011). Spectral data were searched against a target-decoy database consisting of the forward and reverse sequences of the UniProt Danio rerio reference proteome (release 2024_03; 46,844 entries) and a list of common contaminants. Trypsin/P specificity was assigned. A maximum of 2 missed cleavages were tolerated. Carbamidomethylation of cysteine was set as fixed modification. Methionine oxidation and protein N-terminal acetylation were chosen as variable modifications. The “second peptides” option was switched on. The “match between runs” function was activated. The minimum peptide length was set to 7 amino acids. False discovery rate (FDR) was set to 1% at both peptide and protein levels.

The MaxLFQ algorithm (Cox et al., 2014) was employed for label-free protein quantification without using its default normalization option. Minimum LFQ ratio count was set to one. Both the unique and razor peptides were used for quantification. Reverse hits, potential contaminants and “only identified by site” protein groups were filtered out. Normalization of the LFQ protein intensities was performed by median-centring on a logarithmic scale. Proteins were further filtered to retain only those detected in 3 out of the 3 replicates in either group of the comparison. Following imputation of the missing intensity values, a linear model was fitted using the limma package (Phipson et al., 2016) to assess the difference between the two groups for each protein, with adjustment for multiple testing using the Benjamini-Hochberg approach (Benjamini et al., 2001). The log2 fold change and the significance of the difference were displayed on a volcano plot. Only proteins with a minimum log2 fold change of 1 and an FDR-adjusted p value (q value) lower than 0.05 were considered differentially regulated.

### RNA extraction and cDNA preparation

Total RNA was extracted from zebrafish ovaries using TRIzol Reagent (Thermo Fisher Scientific) according to the manufacturer’s instructions. Ovaries were homogenised in 1.5 mL TRIzol using a sterile pestle in RNase-free 1.5 mL microcentrifuge tubes, then centrifuged at 12,000 × g for 5 min at 4°C to remove debris. The supernatant was transferred to a fresh RNase-free tube, and 0.2 mL chloroform per 1 mL TRIzol was added. Samples were vortexed and incubated for 3 min at room temperature before centrifugation at 12,000 × g for 15 min at 4°C. The aqueous phase was transferred to a fresh tube, and RNA was precipitated by adding 0.5 mL isopropanol per 1 mL TRIzol. After 10 min incubation at room temperature, RNA was pelleted by centrifugation at 12,000 × g for 10 min at 4°C. The pellet was washed once with 75% ethanol, air-dried, and resuspended in RNase-free water. RNA concentration and purity were assessed by NanoDrop spectrophotometer. cDNA was synthesised using the ProtoScript II First Strand cDNA Synthesis Kit (New England Biolabs, E6560) according to the manufacturer’s instructions. Sanger sequencing was subsequently performed.

### Primordial germ cell quantification

Primordial germ cells (PGCs) were quantified in zebrafish embryos at 24 hours post fertilisation (hpf) using the Tg(vasa: eGFP) transgenic line, in which PGCs are specifically labelled with GFP. Embryos were manually dechorionated and fixed overnight at 4°C in 4% paraformaldehyde (PFA). Following fixation, embryos were washed in PBS containing 0.1% Tween-20 (PBSTw) and individually transferred to Petri dishes for counting. GFP- positive PGCs were manually counted per embryo under a widefield fluorescence microscope. Total PGC counts per embryo were recorded for subsequent statistical analysis.

### Whole-mount immunofluorescence (4-cell stage embryos)

Whole-mount immunofluorescence was performed on zebrafish embryos at the 4-cell stage. Embryos were fixed for 3 hours at room temperature in 4% PFA with gentle rotation. Following fixation, embryos were washed twice in PBSTw (0.1%) and dehydrated through a methanol series (25%, 50%, 75%, and two steps of 100% methanol; 5 min per step at room temperature). Dehydrated embryos were stored in 100% methanol at −20°C overnight. Prior to immunostaining, embryos were rehydrated through a reverse methanol series back into PBSTw. Antigen retrieval was performed by incubating embryos in retrieval buffer (150 mM Tris-HCl, pH 9.0) for 5 min at room temperature followed by 15 min at 70°C with gentle agitation. Embryos were blocked in 8% bovine serum albumin (BSA) in PBSTw for 1 h at room temperature. Primary antibodies (1:250) were diluted in 8% BSA/PBSTw and incubated overnight at 4°C with gentle rotation. Embryos were washed six times for 30-60’ each time in PBSTw at room temperature. Secondary antibodies (1:500) diluted in 8% BSA/PBSTw were added, and embryos were incubated overnight at 4°C in the dark with gentle rotation. Following secondary incubation, embryos were washed twice in PBSTw (30 min per wash), then mounted in imaging chambers (Ibidi μ-slide 8-well) for fluorescence microscopy.

### Whole-mount immunofluorescence (oocytes)

Whole-mount immunofluorescence was performed on isolated zebrafish oocytes. Oocytes were fixed for 1 hour at room temperature in 2% PFA with gentle rotation. Following fixation, oocytes were washed twice in PBS Triton (0.3%) and dehydrated through a methanol series (as above). Dehydrated oocytes were stored in 100% methanol at −20°C overnight for a minimum of one night. Prior to immunostaining, oocytes were rehydrated through a reverse methanol series back into PBS Triton. Antigen retrieval was performed by incubating oocytes in retrieval buffer (150 mM Tris-HCl, pH 9.0) for 5 min at room temperature followed by 15 min at 70°C with gentle agitation. Oocytes were blocked in 8% bovine serum albumin (BSA) in PBSTw for 1 h at room temperature. Primary antibodies (1:100) were diluted in 8% BSA/PBSTw and incubated overnight at 4°C with gentle rotation. Next day, oocytes were washed six times over 1 h in PBSTw at room temperature. Secondary antibodies (1:500) diluted in 8% BSA/PBSTw were added, and oocytes were incubated overnight at 4°C in the dark with gentle rotation. Following secondary incubation, oocytes were washed twice in PBSTw (30 min per wash), then mounted in imaging chambers (Ibidi μ-slide 8-well) for fluorescence microscopy.

### Whole-mount immunofluorescence (24 hpf embryos)

Whole-mount immunofluorescence was performed on zebrafish embryos at 24 hpf. Embryos were manually dechorionated using forceps and transferred to 2 mL microcentrifuge tubes. Embryos were fixed overnight at 4°C in 4% PFA with gentle rotation.

Following fixation, embryos were washed twice in PBSTw (0.1%) and dehydrated through a methanol series (as above). Dehydrated embryos were stored in 100% methanol at −20°C for at least one overnight. Prior to immunostaining, embryos were rehydrated through a reverse methanol series back into PBSTw.

Antigen retrieval was performed as above. Embryos were then blocked in 8% bovine serum albumin (BSA) in PBSTw for 1 h at room temperature. Primary antibodies (1:250) were diluted in 8% BSA/PBSTw and incubated overnight at 4°C with gentle rotation. Embryos were subsequently washed six times over 1 h in PBSTw at room temperature. Secondary antibodies (1:500) diluted in 8% BSA/PBSTw were added, and embryos were incubated overnight at 4°C in the dark with gentle rotation. Following secondary incubation, embryos were washed twice in PBSTw (30 min per wash), then mounted in imaging chambers (Ibidi μ-slide 8-well) for fluorescence microscopy.

### Single-molecule fluorescence

Single-molecule FISH (smFISH) was performed on whole-mount zebrafish embryos at the 4-cell stage, optionally combined with immunofluorescence. Fixed embryos were incubated in hybridisation buffer (10% dextran sulfate, 10% formamide, 1 mg/mL tRNA, 2× SSC, 0.02% BSA, and 2 mM vanadyl-ribonucleoside complex) for 1 h at 30°C.

Probe sets were added at 1:100 dilution in hybridisation buffer and incubated overnight at 30°C. For combined smFISH/immunofluorescence, primary antibody (1:100 for ovaries, 1:250 for embryos) was added at this step. The following day, samples were washed once for 15 min at 30°C with wash buffer. A second wash with added secondary antibody (1:500) was conducted for 30’. Then a further two washes at 15’ each. Samples were mounted in ibidi Mounting Medium (ibidi, cat. no. 50001) in µ-Slide 8 Well chambers (ibidi, Cat.No:80806).

### Recombinant protein expression and purification

His14-TEVsite-tagged Buc/Buc2l fragments were expressed from pQE plasmids in E. coli C41-RIL. Cells were grown in 4 to 8 L TB medium at 37°C until an OD (600) of 1 and expression was induced using 0.1 mM IPTG at 37°C for 3 hours. Cells were harvested (4k x g for 15 min at 20°C), lysed by sonication in 70-140 ml lysis buffer (50 mM Tris-HCl, 500 mM NaCl, 6 M GdnHCl, 10 mM imidazole, 0.1% Triton X-100, 10 mM 2- Mercaptoethanol, 1 mM EDTA, pH 8.0) and cleared using centrifugation (55k x g, 30 min, 20°C). Proteins were enriched from cleared lysate using 10 ml Ni-INDIGO agarose slurry in batch for 3 h at room temperature. Beads were washed extensively (50 mM Tris-HCl, 500 mM NaCl, 4 M GdnHCl, 10 mM imidazole, 10 mM 2-Mercaptoethanol, pH 8.0) before elution in 20 ml (50 mM Tris-HCl, 500 mM NaCl, 4 M GdnHCl, 500 mM imidazole, 10 mM 2-Mercaptoethanol, pH 8.0), followed by spin-concentration and size exclusion chromatography (Superdex 200 16/60 in SEC buffer: 20 mM Hepes-NaOH, 4 M GdnHCl, 1 mM TCEP, pH 7.4). Purified proteins were spin-concentrated, aliquoted, snap-frozen in liquid nitrogen and stored at −80°C.

Purified recombinant proteins were incubated at 120 µM with 2.5-fold molar excess of maleimide dyes (Atto647-maleimide, Atto565-maleimide or Atto488-maleimide) at room temperature for 1.5–2.5 hours in SEC buffer and a total volume of 500 µl. Unreacted dye was quenched by addition of 5 mM DTT and samples were centrifuged (20k x g, 5 min, 20°C). Free dye and DTT were removed using NAP-5 desalting columns in SEC buffer), followed by spin-concentration aliquoting, snap-freezing in liquid nitrogen and storage at −80°C.

### In vitro phase separation assays

Recombinant proteins were subjected to phase separation by dilution-induced removal of guanidinium chloride. Labelled and unlabelled proteins were mixed at final concentrations of 100–2000 nM with 5% labelled protein in phase separation buffer (20 mM HEPES- NaOH pH 7.4, 150 mM NaCl, 1 mM DTT) (Fig. S5A). For the salt series, proteins were diluted to 500 nM in the same buffer containing 0, 50, 150 or 300 mM NaCl (Fig. S5B, C). Phase separation was induced by diluting proteins from storage buffer (4 M guanidinium chloride) into phase separation buffer, reducing the denaturant concentration to 20 mM guanidinium chloride for single-protein assays and 40 mM for assays in which two proteins were mixed. Samples were incubated at room temperature for 10 minutes to allow condensate formation, then transferred to ibidi μ-Slide 15 Well 3D imaging chambers, and left for 5 minutes. Condensates were imaged on a spinning disk confocal microscope. Z- stacks were acquired and analysed using Imaris (Bitplane) to quantify condensate volume, number and morphology.

### Imaging and microscopy

Fluorescence imaging was performed using multiple microscope systems depending on the application. Widefield fluorescence microscopy was used for PGC quantification and initial immunofluorescence screening. For higher-resolution imaging, confocal microscopy was performed using a Visitron System spinning disk confocal microscope to image embryos and oocytes. For smFISH in embryos, the BC43 Spinning Disk Confocal microscope was used. Super-resolution imaging of germ plasm was performed using Stellaris 8 FALCON confocal and additional imaging conducted using Leica TCS SP5 STED CW microscope. Images were processed using Fiji (ImageJ) and Imaris software (Bitplane).

### Assembly segmentation and quantification

Confocal z-stacks were analysed in Imaris (Oxford Instruments, version 11) using the Surfaces module. As assembly size and fluorescence intensity varied substantially within and between images, precluding accurate segmentation of both large and small assemblies with a single intensity threshold, assemblies were segmented in two size- stratified passes.

In the first pass, an intensity threshold was set to capture large, bright assemblies only, validated by visual comparison of the rendered surface against the raw channel. In the second pass, a lower intensity threshold was applied to capture the smaller, dim assemblies. To prevent duplicate counting of assemblies detected in both passes, objects in the small-assembly population that spatially overlapped with objects in the large- assembly surface were identified with overlapped ratio statistics and then excluded from the small-assembly population. Deduplication was applied within each protein channel only, so that colocalization between the two proteins was unaffected. The resulting large and small assembly populations were exported as separate statistics tables and combined for quantification of assembly number, volume, and mean intensity. Five to thirteen fields of view were acquired per well. Objects below 0.04 µm³ (approximately 17 voxels at the binned voxel size) were excluded as segmentation noise; this threshold removed less than 2% of total condensed volume. Sphericity was scored only on objects of at least 0.1 µm³ that did not touch an image border. Assemblies touching the image border were retained for number and volume measurements, as these represent genuine, partially imaged assemblies; a sensitivity analysis excluding assemblies cut by the top of the z-stack gave equivalent results. Colocalization was quantified per field as the percentage of one protein’s total assembly volume that overlapped assemblies of the partner protein, calculated from Imaris overlapped volume ratio statistics. Data handling and plotting were performed in R (version 4.4) using tidyverse, emmeans and ggplot2.

### Statistics - In vitro assembly assays

*In vitro* assembly assays. Values were measured per field of view and averaged within each well, so that the unit of analysis was the independent experiment (n = 3) rather than the individual field or assembly. Fields containing no assemblies counted as zero for assembly number and total volume. Mean assembly size was only calculated for wells in which most fields (>50%) contained at least three assemblies; other wells are reported as not determined (n.d.).

Assembly number, total volume and mean size were log10-transformed, and constructs were compared using a linear model with replicate as a blocking factor. Each construct was compared with full-length Buc2l, and p-values were Holm-corrected. To test whether assembly size depended on A-unit number, the same model was used with A-unit number as a continuous predictor (full-length Buc2l, Δ5, Δ3-5, Δ2-5 and Δ1-5 = 5, 4, 2, 1 and 0 A- units); the reported p-value is that of the A-unit term. Proteins assayed alone and in mixtures came from separate experiments and were therefore compared using unpaired Welch’s t-tests, reported uncorrected. Effect sizes are given as fold changes.

---

**Supplementary Figure 1.**
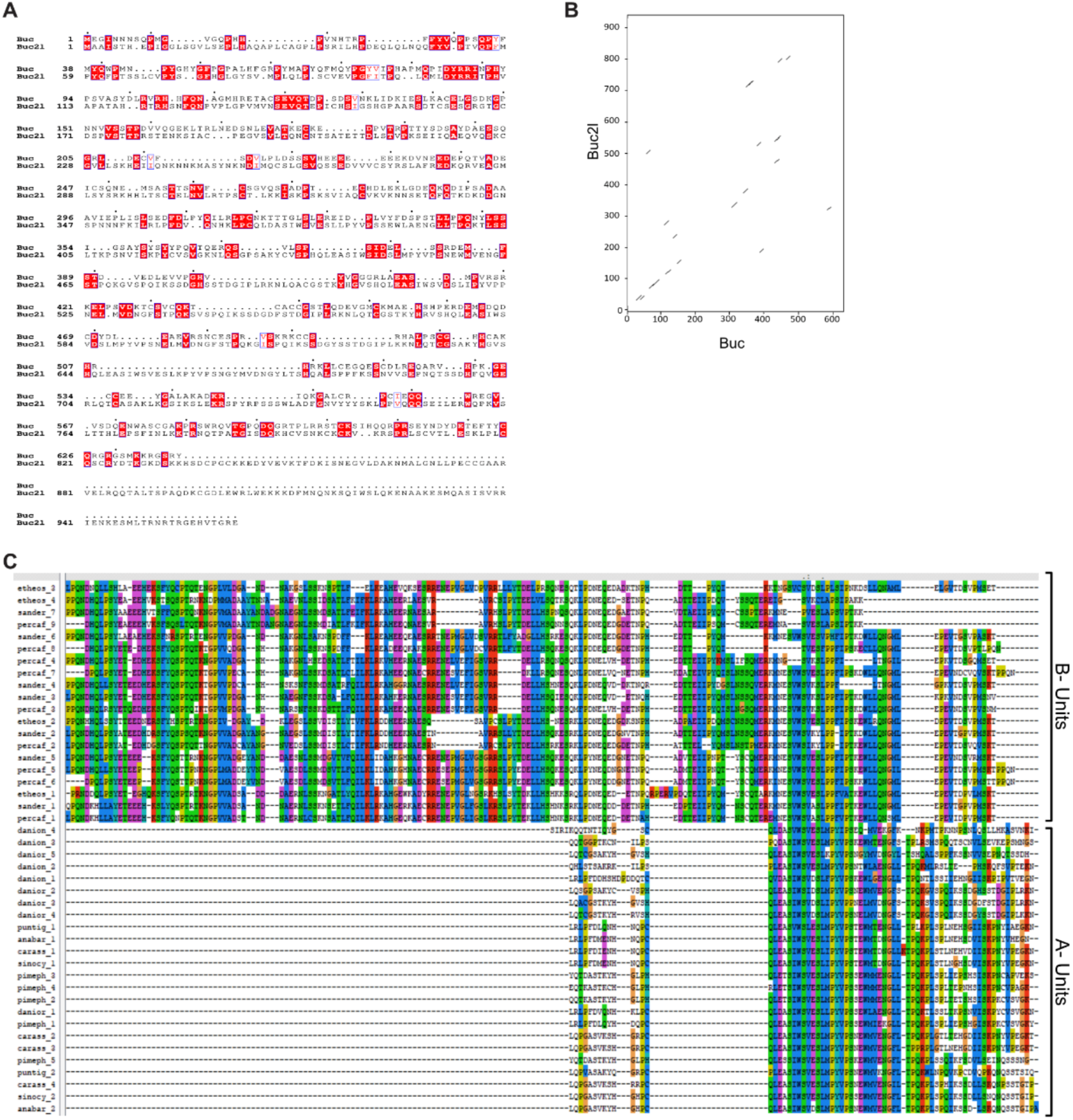
Sequence comparison of Buc and Buc2l, and alignment of Buc2l repeat units. (A) Pairwise alignment of zebrafish Buc and Buc2l (EMBOSS Needle; 18.2% identity, 28.4% similarity). Identical and similar residues are indicated. (B) Dot plot comparing zebrafish Buie and Buc2l; similarity is concentrated in the N-terminal ∼200 residues. (C) Multiple sequence alignment of individual Buc2l repeat units from cyprinids (A-units) and percids (B-units). Sequence names indicate species and repeat number.

**Supplementary Figure 2.**
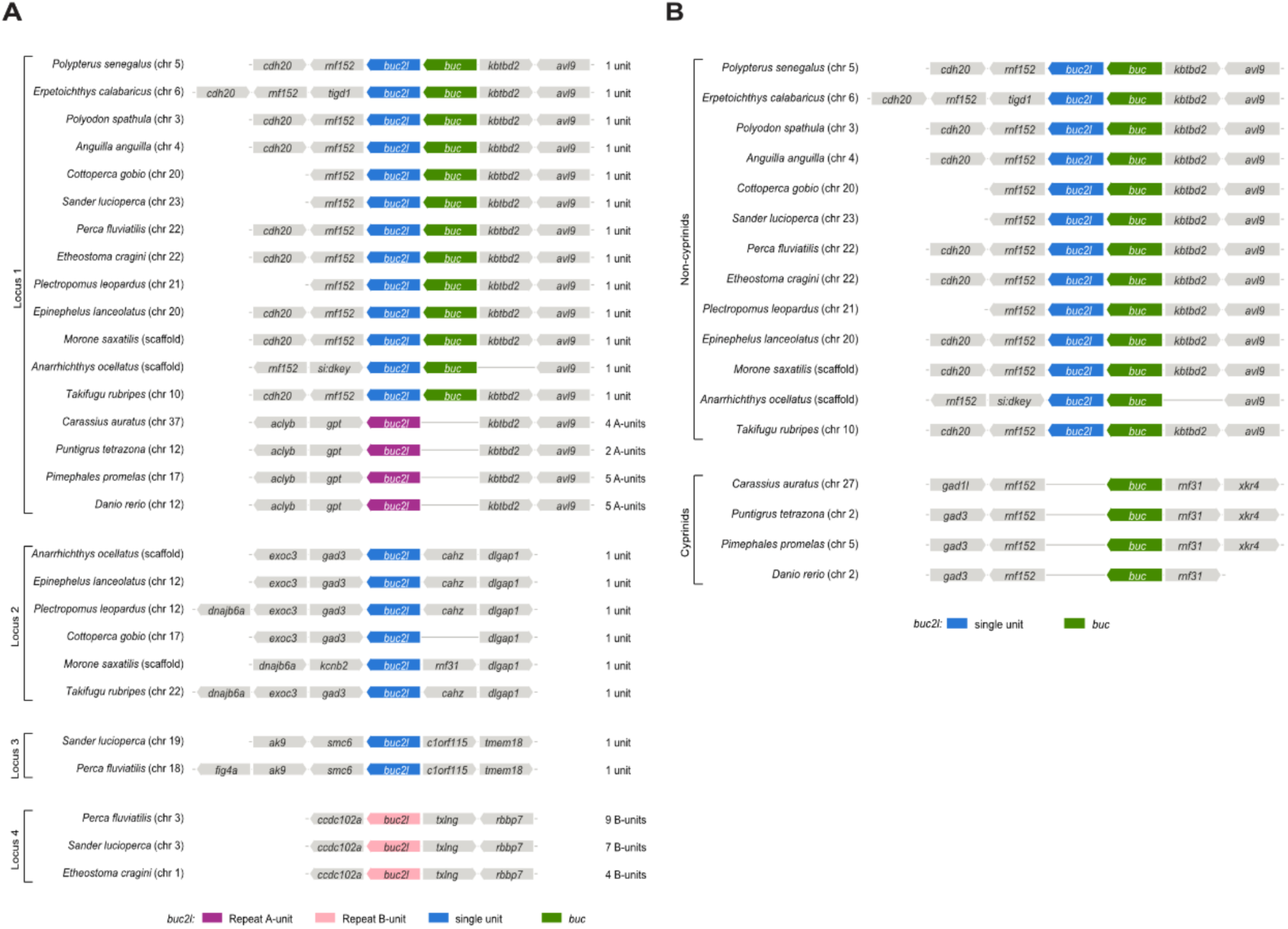
Synteny of *buc2l* and *buc* loci across fish. (A) Gene order around each gene, grouped by locus (Loci 1-4). *buc2l* genes are coloured by repeat type (A-units, B-units or single unit), and the *buc2l* number of repeat units is given on the right. Lines indicate the absence of a gene. In cyprinids, *buc* is absent from Locus 1 and is found on a separate chromosome (see B). (B) Genomic context of *buc* in non-cyprinid and cyprinid fish. In non- cyprinids, *buc* lies directly next to *buc2l.* In cyprinids, *buc2l* is absent from this locus, but *buc* retains its neighbour *rnf152*

**Supplementary Figure 3.**
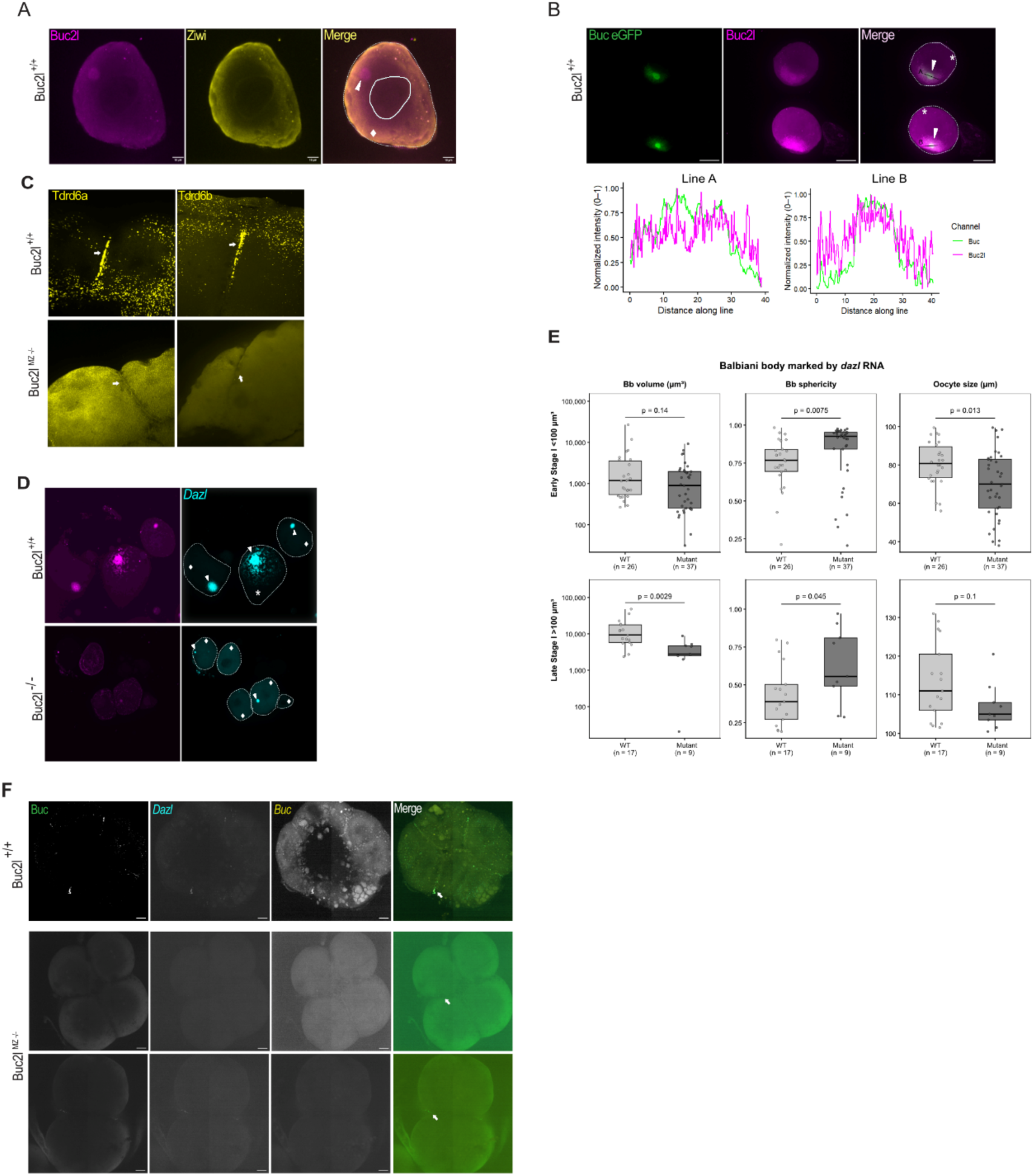
Buc2l localisation and Balbiani body and germplasm defects in buc2l mutants. (A) Wild-type oocyte immunostained for Buc2l and Ziwi. The oocyte and nucleus are outlined, and the Balbiani body is indicated. (B) Oocytes from *Tg(buc.buc-EGFP)* ovaries immunostained for Buc2l, with normalised intensity profiles of Buc-EGFP and Buc2l along lines A and B (below). (C) Tdrd6a and Tdrdβb immunostaining at the cleavage furrow of 4-cell stage wild-type and buc2/^MZ-/-^ embryos. Germplasm is indicated. (D) Wild-type and *buc2/^-/-^* oocytes immunostained for Buc2l and labelled for *dazl* RNA by smFISH. Oocytes are outlined, Balbiani bodies and oocyte stages are indicated. (E) Balbiani body volume, Balbiani body sphericity and oocyte size in early (<100 μm; top) and late (>100 μm; bottom) stage I oocytes from wild-type and *buc2l^-/-^* females, with the Balbiani body marked by dazl RNA. Each point is one oocyte. Genotypes were compared by two-sided Mann-Whitney U test, Holm-corrected. (F) Buc protein, *dazl* RNA and *buc* RNA in 4-cell stage wild-type and *buc2l^MZ-/-^*embryos (two representative mutant embryos shown). Germplasm is indicated. Oocytes are outlined, and oocyte stages are indicated, early (diamond) and late (asterisk). Scale bars, 10 μm (A), 20 μm (B), 50 μm (F).

**Supplementary Figure 4.**
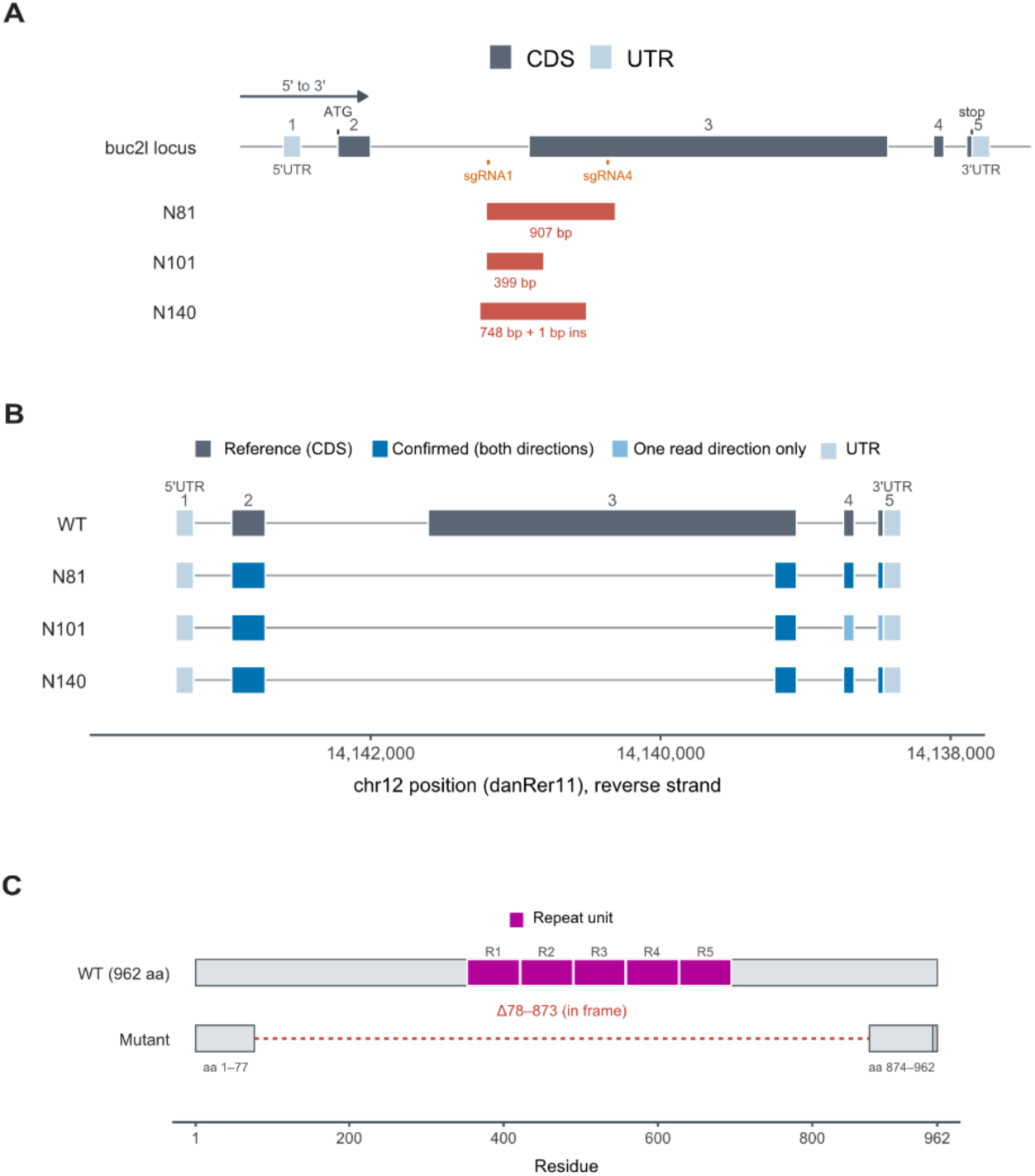
Generation and characterisation of buc2l mutant alleles. (A) The *buc2l* locus, showing the sgRNAI and sgRNA4 target sites in exon 3 and the genomic lesions in the N81 (907 bp deletion), N101 (399 bp deletion) and N140 (748 bp deletion with 1 bp insertion) alleles. (B) *buc2l* transcript structure in wild type and in the N81, N101 and N140 alleles, determined by RT-PCR and Sanger sequencing of cDNA and mapped to chromosome 12 (danReríl, reverse strand). In all three alleles, exon 2 is spliced directly to the 3’ end of exon 3, so most of exon 3 is missing from the transcript. (C) Predicted Buc2l protein in wild type (962 aa) and in the N81, N101 and N140 alleles. All three encode the same in-frame deletion of residues, which removes the entire repeat array (R1-R5)

**Supplementary Figure 5.**
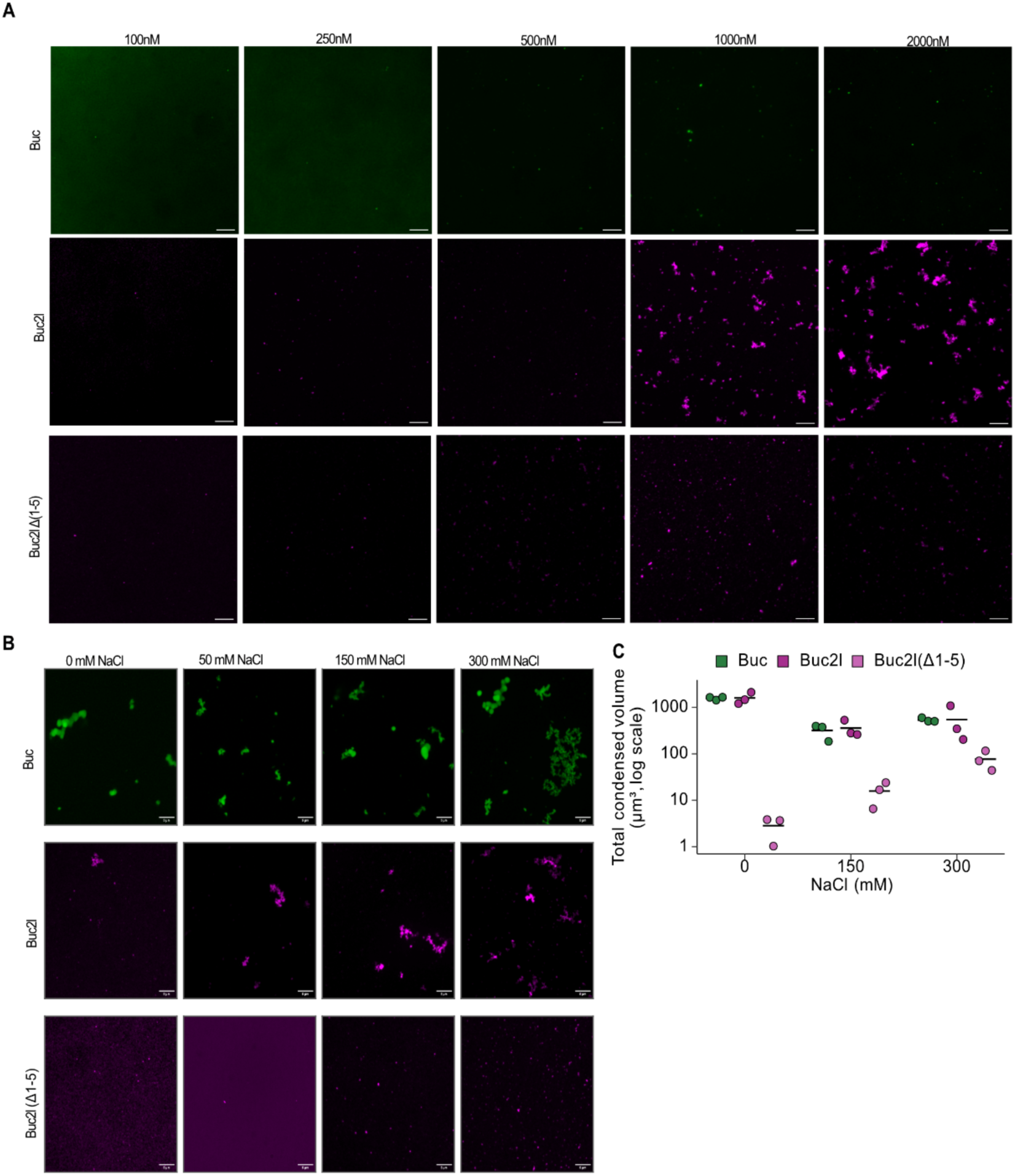
Concentration and salt dependence of Buc, Buc2l and Buc2l(Δ1-5) assembly. (A) Buc (green), Buc2l and Buc2l(Δ1-5) (magenta) alone at 100-2000 nM, imaged 15 min after mixing, representative images from a single experiment. (B) Buc, Buc2l and Buc2l(Δ1-5) (500 **nM)** at 0-300 mM NaCI. (C) Total condensed volume per field (log scale) at 0, 150 and 300 mM NaCI; dots, individual fields from a single experiment; lines, mean, no statistical comparison was made. Maximum-intensity projections. Scale bars, 10 μm (A), 5 μm (B).

**Supplementary Figure 6.**
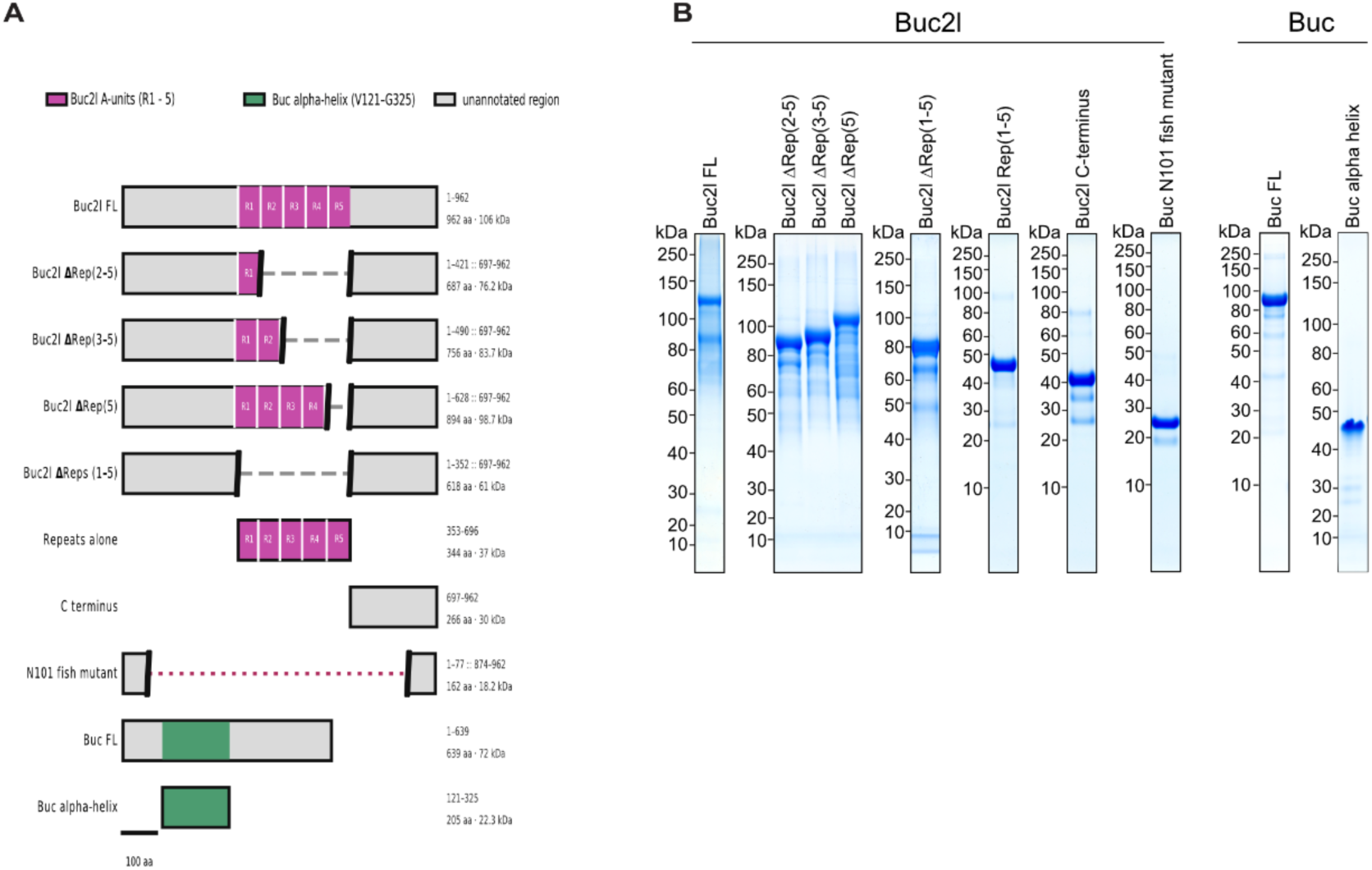
Recombinant Buc and Buc2l proteins used in this study. Schematic of full-length Buc2l (FL), A-unit deletions (Δ5, Δ3-5, Δ2-5, Δ1-5), the A-unit region alone (R1-5), the C- terminus (697-962), the protein encoded by the buc2l^+^ allele (N101; Δ78-873), full-length Buc and the Buc α-helix fragment (121-325). Residues retained and predicted molecular weight are given at right. Scale bar, 100 aa. (B) Coomassie-stained SDS-PAGE of recombinant proteins used in this study

**Supplementary Figure 7.**
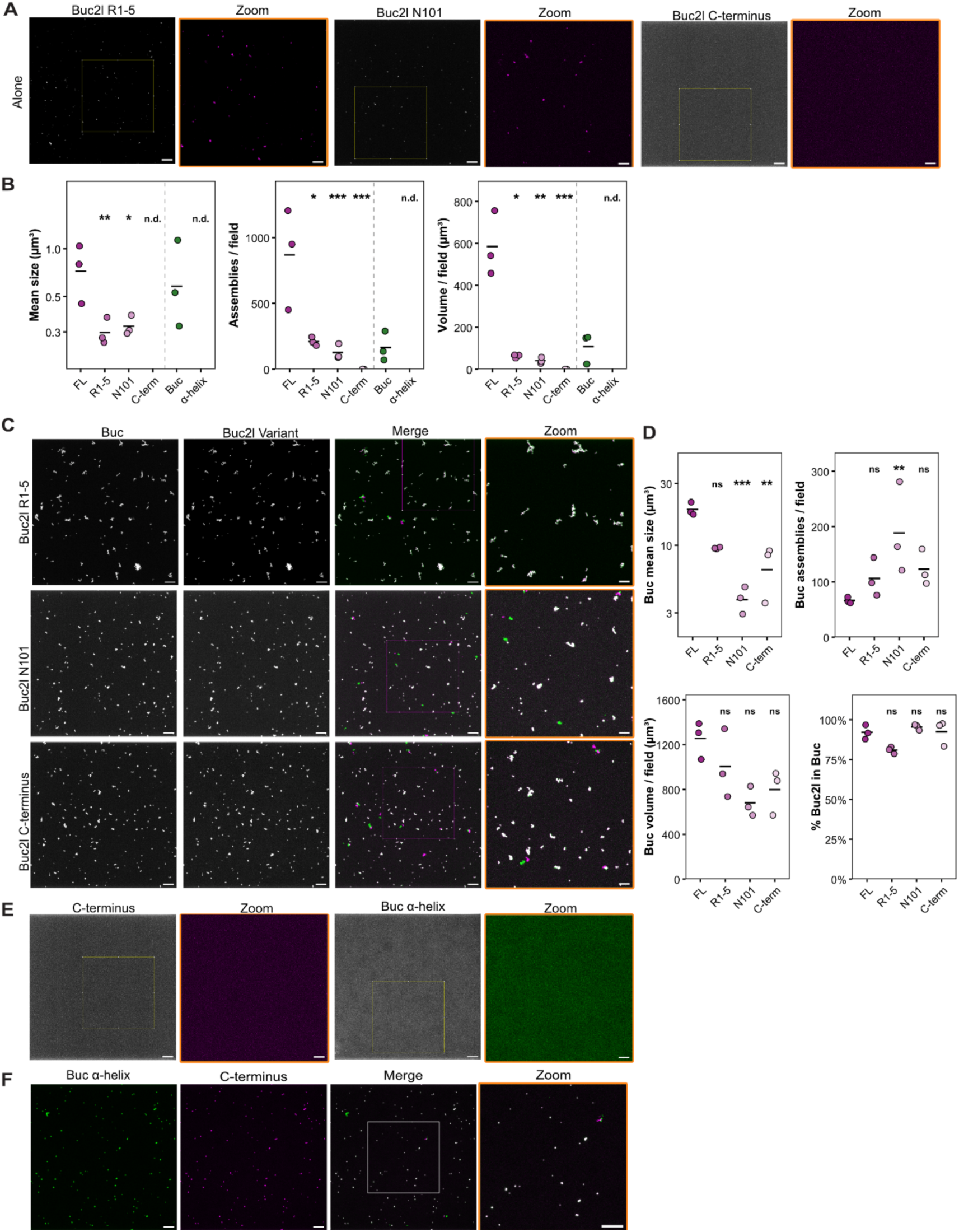
Buc2l fragments and the Buc α-helix form few or no assemblies on their own but are recruited into Buc assemblies. (A) Buc2l(R1-5), Buc2l N101 and Buc2l C-terminus alone (500 nM); zooms of boxed regions on the right. (B) Quantification of proteins alone (A, E); fragments vs FL, α-helix vs Buc. (C) Buc combined with each Buc2l fragment (500 nM each): Buc, Buc2l variant, merge (Buc, green; Buc2l, magenta) and zoom. (D) Quantification of C (Buc channel) and % Buc2l- variant in Buc assemblies; vs Buc + FL. (E) Buc2l C-terminus and Buc α-helix alone (500 nM), zooms of boxed regions. (F) Buc α-helix combined with Buc2l C-terminus (500 nM each): α-helix, C-terminus, merge and zoom. Statistics as in Figure 7; Buc α-helix was compared with Buc rather than with Buc2l. Scale bars, 10 μm; zooms, 5 μm. Dots, individual replicates (mean of all fields per well); bars, mean of replicates; n = 3 (C-terminus alone, n = 2). *p < 0.05, **p < 0.01, ***p < 0.001; ns, not significant; n.d., not determined (no assemblies formed).

